# A-to-I RNA editing in honeybees shows signals of adaptation and convergent evolution

**DOI:** 10.1101/2020.01.15.907287

**Authors:** Yuange Duan, Shengqian Dou, Hagit T. Porath, Jiaxing Huang, Eli Eisenberg, Jian Lu

## Abstract

Social insects exhibit extensive phenotypic diversities among the genetically similar individuals, suggesting a role for the epigenetic regulations beyond the genome level. The ADAR-mediated adenosine-to-inosine (A-to-I) RNA editing, an evolutionarily conserved mechanism, facilitates adaptive evolution by expanding proteomic diversities. Here, we characterize the A-to-I RNA editome of honeybees (*Apis mellifera*), identifying 407 high-confidence A-to-I editing sites. Editing is most abundant in the heads, and shows signatures for positive selection. Editing behavior differs between foragers and nurses, suggesting a role for editing in caste differentiation. Although only five sites are conserved between bees and flies, an unexpectedly large number of genes exhibit editing in both species, albeit at different locations, including the nonsynonymous auto-editing of *Adar*. This convergent evolution, where the same target genes independently acquire recoding events in distant diverged clades, together with the signals of adaptation observed in honeybees alone, further supports the notion of recoding being adaptive.

## INTRODUCTION

Adenosine (A) to inosine (A-to-I) RNA editing, catalyzed by enzymes of the ADAR (adenosine deaminase acting on RNA) family (Savva et al. 2012b), is an evolutionarily conserved mechanism that expands RNA diversity at the co-transcriptional or post-transcriptional level in metazoans (Bass 2002; Nishikura 2010; Eisenberg and Levanon 2018). Two catalytically-active *Adar* genes are encoded in most metazoans. However, insects have lost ADAR1 and encode only a single *Adar* gene (Keegan et al. 2011). Due to the structural similarity between inosine (I) and guanosine (G), I is generally believed to be recognized as G in many cellular processes such as mRNA splicing (Rueter et al. 1999; Flomen et al. 2004; Jin et al. 2007; Lev-Maor et al. 2007), microRNA (miRNA) biogenesis or target recognition (Liang and Landweber 2007; Borchert et al. 2009; Alon et al. 2012), and mRNA translation (Basilio et al. 1962; Licht et al. 2019). Therefore, an A-to-I RNA editing usually has a similar effect as an A-to-G DNA substitution. A-to-I editing plays essential roles in many biological and physiological processes(Keegan et al. 2001; Nishikura 2010), and dysregulation of A-to-I editing might be associated with cancer, autoimmune disorders, or other human diseases (Gallo et al. 2017).

During the past two decades A-to-I RNA editing sites have been systematically characterized in various metazoan species (Ramaswami and Li 2016). The majority of RNA editing sites are located in clusters in non-coding regions of humans (Athanasiadis et al. 2004; Blow et al. 2004; Kim et al. 2004; Levanon et al. 2004; Picardi et al. 2017), monkeys (Chen et al. 2014; Yang et al. 2015), mice (Neeman et al. 2006; Danecek et al. 2012), worms (Morse and Bass 1999; Zhao et al. 2015; Goldstein et al. 2017), corals (Porath et al. 2017b) and many other species (Porath et al. 2017a). In most species, only a minute fraction of the edits resides within the coding sequence. Notable exceptions are *Drosophila* (Graveley et al. 2011; Rodriguez et al. 2012; St Laurent et al. 2013; Mazloomian and Meyer 2015; Yu et al. 2016; Buchumenski et al. 2017; Duan et al. 2017; Zhang et al. 2017) and cephalopod species (Alon et al. 2015; Liscovitch-Brauer et al. 2017). Despite the deep conservation of A-to-I editing mechanism, the target landscapes of editing have considerably evolved during metazoan evolution. Only one editing site is known to be conserved across virtually all mammals, *Drosophila*, and cephalopods (Porath et al. 2019), and it thus seems that RNA editing modulates the diversity of the transcriptomes and proteomes in a lineage-specific manner.

The forces driving the evolution of A-to-I editing across species at the macro-evolutional scale are not well understood. RNA editing was hypothesized to facilitate adaptive evolution by increasing proteomic diversities temporally or spatially, in a manner more flexible than genomic mutations (Gommans et al. 2009; Nishikura 2010; Klironomos et al. 2013; Rosenthal 2015; Nishikura 2016). However, in most species studied the ratio of nonsynonymous *(N)* to synonymous (*S*) editing sites (*N/S*) is lower than expected for random sites, suggesting that recoding events may be overall non-adaptive (Xu and Zhang 2014). Here too *Drosophila* and cephalopods stand out as exceptions, exhibiting high *N/S* ratios indicating positive selection of recoding. Furthermore, hundreds of recoding sites were shown to be conserved across the *Drosophila* lineage (Graveley et al. 2011; Rodriguez et al. 2012; St Laurent et al. 2013; Mazloomian and Meyer 2015; Yu et al. 2016; Buchumenski et al. 2017; Duan et al. 2017; Zhang et al. 2017), and thousands in the behaviorally sophisticated cephalopods (Garrett and Rosenthal 2012; Alon et al. 2015; Liscovitch-Brauer et al. 2017), supporting the functional importance of these editing events. However, in only a few examples was the advantageous effect conferred by RNA editing explicitly demonstrated (see for example (Garrett and Rosenthal 2012)), and even it is not yet clear to what extent editing is indeed utilized for proteome diversification.

Social insects, including bees and ants, show extensive phenotypic plasticity. The morphologically and behaviorally differentiated social castes such as queens, workers, and drones have the same set of diploid or haploid genome. The social insects provide us with model system to study how phenotypic diversity is regulated (Page et al. 2012; Yan et al. 2014). RNA editing is well suited to contribute to the behavioral variation among genetically similar individuals. Indeed, differential editing was demonstrated for a few sites in leaf-cutting ants and in worker bumblebees (Li et al. 2014; Porath et al. 2019).

Here we wish to study the contribution of RNA editing to caste differentiation in honeybee (*Apis mellifera*), an important pollinator and a model for complicated social behaviors in insects (Page et al. 2012). Unlike bumblebees, honeybees have a sharp task-specialization, with distinct worker castes. Genetic mapping demonstrated that the phenotypic plasticity in honeybees is associated with complex epistatic and pleiotropic genetic networks that influence reproductive regulation and foraging behaviors (Page et al. 2012). We wish to check whether A-to-I RNA editing may contribute to the proteomic diversity underlying this phenotypic plasticity. We investigate the A-to-I RNA editome in different tissues of four honeybee drone individuals and detect over four hundred A-to-I sites in multiple tissues. Editing is enriched in the head, and exhibit signs for positive selection and a particularly high *N/S* ratio. We show editing is elevated in foragers compared to nurses. Five editing sites are conserved between honeybee, bumblebee and *Drosophila*. Interestingly, we find a significantly high number of cases where the same gene in edited in both bees and flies, even if not at the same position. One example is the auto-editing of *Adar* mRNA in bees and flies, which might play an auto-regulatory role in the two clades. This finding may further support possible convergent evolution and adaptation.

## RESULTS

### Editome of honeybee

We deep-sequenced genomic DNA and RNA from the head, thorax, and abdomen for each of four individual drones (12 RNA-seq samples) (Figure 1A). The transcriptome libraries of drones 1 and 2 were constructed by selecting polyA tailed mRNAs while the libraries of drones 3 and 4 were constructed by using the Ribo-Zero kit to deplete ribosomal RNAs (Transparent Methods). The male drone is haploid, simplifying identification of its genomic SNPs. DNA-Seq reads were mapped to the reference genome to identify 826,890-858,949 SNPs in each of the four drones, and 1,303,225 unique sites combined (Transparent Methods; Table S1). Median sequencing coverage at the SNP sites was 39, and in 96.6% of the sites all DNA reads supported the SNP (Supplemental Fig S1). The identified SNPs were then used to produce a masked genome version where the reference allele is substituted by the individual-specific alleles identified for each drone, to facilitate accurate detection of RNA editing sites (Figure 1B; Transparent Methods).

**Figure 1.**
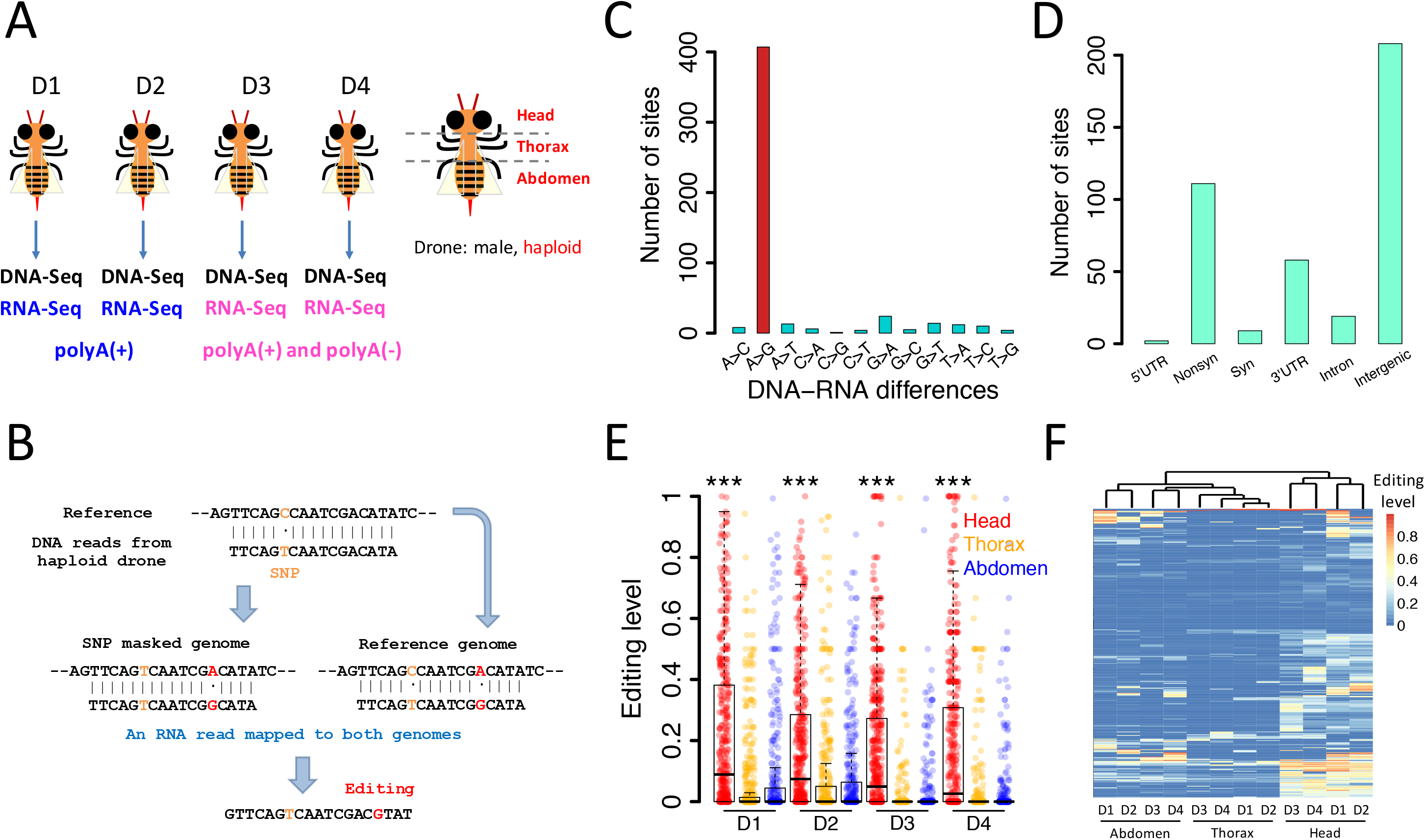
Identification and annotation of the A-to-I RNA editing sites in honeybee. (A) Workflow of sample collection, dissection and library construction. (B) Identification of A-to-I RNA editing sites is facilitated by mapping to both the reference genome and the masked genome. (C) Distribution of DNA-RNA mismatch types detected in honeybee. (D) Distribution of the detected A-to-I editing sites over different genic regions. (E) Editing levels at the detected as measured in three tissues of each of four individuals (D1-D4). Editing levels are higher in the head (head versus pooled thorax and abdomen, Wilcoxon rank sum test; ***: *P* < 0.001. Exact *P* values, left to right: 2.1e-35, 5.7e-20, 4.0e-29, and 3.9e-38). (F) The editing profiles for the twelve tissues cluster according to their tissue of origin.

For each of the 12 RNA-Seq samples, 9.2-21.5 million reads were uniquely mapped to the reference genome, and similar numbers of reads were mapped to the masked genome sequences (Transparent Methods; Table S2). Analyzing pooled data for each of the three tissues separately, we identified (FDR=0.05) 376 (84.3%) A-to-G sites among 446 variations in heads, 106 (66.7%) A-to-G sites among 159 variations in thoraxes, and 137 (67.5%) A-to-G sites among 203 variations in abdomens (Figure S2). Combined, we obtained 407 (80.1%) unique A-to-G sites among 508 variations (Figure 1C). These 407 sites are regarded as A-to-I RNA editing sites (Table S3, Information of the candidate editing sites identified in this study, Related to Figure 1). The nucleotide context around these putative editing sites is consistent with the known ADAR binding motif (Figure S3). The median DNA coverage (over the 407 A-to-I editing sites) was 25 reads, sufficient to exclude possible SNPs (Figure S4).

To improve mapping accuracy, we discarded variants in repeat regions, and required a variation site to be found by mapping to both the reference genome and the masked genome (Transparent Methods). To test whether these criteria are too stringent, we checked the variation sites discarded by the two filtering steps. First, we looked at the variation sites located in the repeat regions. Without any filter, we found 19,573 unique variation sites that overlapped with repeat regions, only 3,694 (19%) of which were A-to-G variants. Following a binomial test and multiple testing correction (to remove random sequencing errors), 1,277 variation sites were maintained, only 324 (25%) of them A-to-G variations. Thus, filtering repetitive regions does contribute appreciably to the precision of our detection. Consistently, hyper-editing analysis did not reveal multiple sites in repetitive regions (Transparent Methods).

As for the masked genome filter, only 38% (98/256) of the variation sites found by mapping to the masked genome but not to the reference genome are A-to-G mismatches. Following binomial test for sequencing error and multiple testing correction for *P* values, only 41 variation sites were maintained, none of them was an A-to-G variation. Conversely, only 31% (82/268) variation sites found by mapping to the reference genome but not to the masked genome are A-to-G mismatches. Of these 268 variation sites, 82 were maintained after multiple testing correction, 20 of them were A-to-G variations. Thus, most of the sites supported by only one method of mapping are likely not due to A-to-I RNA editing.

Among the 407 editing sites we identified, 199 sites are located in gene regions, including 111 nonsynonymous (*N*) sites, 9 synonymous (*S*) sites, and several sites in UTRs and introns (Figure 1D). The other 208 sites are annotated as intergenic. As expected from the different library construction strategies, the fraction of sites in coding regions is higher and the intergenic fraction is lower for sites observed in drones 1 and 2 compared with drones 3 and 4 (23.2 ± 2.6% in CDS in drones 1 and 2 versus 14.5 ± 5.1% in CDS in drones 3 and 4; 54.9 ± 2.2% in intergenic regions in drones 1 and 2 versus 70.5 ± 5.4% in intergenic regions in drones 3 and 4). The number of sites for which editing is observed in each sample (per-sample editing level > 0) varies considerably (40-278 sites per sample; Table S4). The partial overlap between these sets of sites is mostly due to sites being undetected at a given sample due to low per-sample coverage (Figure S5).

Interestingly, editing seems to be enriched in heads. The number of detected sites (Figure S6), the editing levels (Figure 1E) and the fraction of coding sites (Figure S7), are all higher in heads. Clustering the samples by their editing profile, the three tissue-types form distinct clusters (Figure 1F). Consistently, *Adar* expression is also higher in heads than other tissues (Figure S8). Looking at specific sites, 78 sites (out of 378 sites with sufficient coverage, see Transparent Methods) are differentially-edited in the head, compared to non-head tissues. In all of these 78 sites, editing is higher in the head (Table S3, Information of the candidate editing sites identified in this study, Related to Figure 1).

### Signals of adaptation

The nonsynonymous to synonymous (*N*/*S*) ratio for sites in the coding region is 111/9 = 12.3 (Table S4), compared to a ratio of 2.26 observed for random adenosine to guanosine substitutions (see Transparent Methods). This strongly suggests that the nonsynonymous editing events in honeybees are overall adaptive. Notably, the *N*/*S* is exceptionally high, even compared to the ratios we previously found in the brains of *Drosophila* (Duan et al. 2017) or in cephalopods (Alon et al. 2015; Liscovitch-Brauer et al. 2017). Looking at each sample separately, at pooled tissue data, or focusing on sites that appear in at least two individuals per tissue all show the *N*/*S* ratio to be especially high in heads (Figure 2A and Table S4). Furthermore, the editing level at nonsynonymous sites is significantly higher than synonymous sites editing levels in heads of honeybees (Figure 2B).

**Figure 2.**
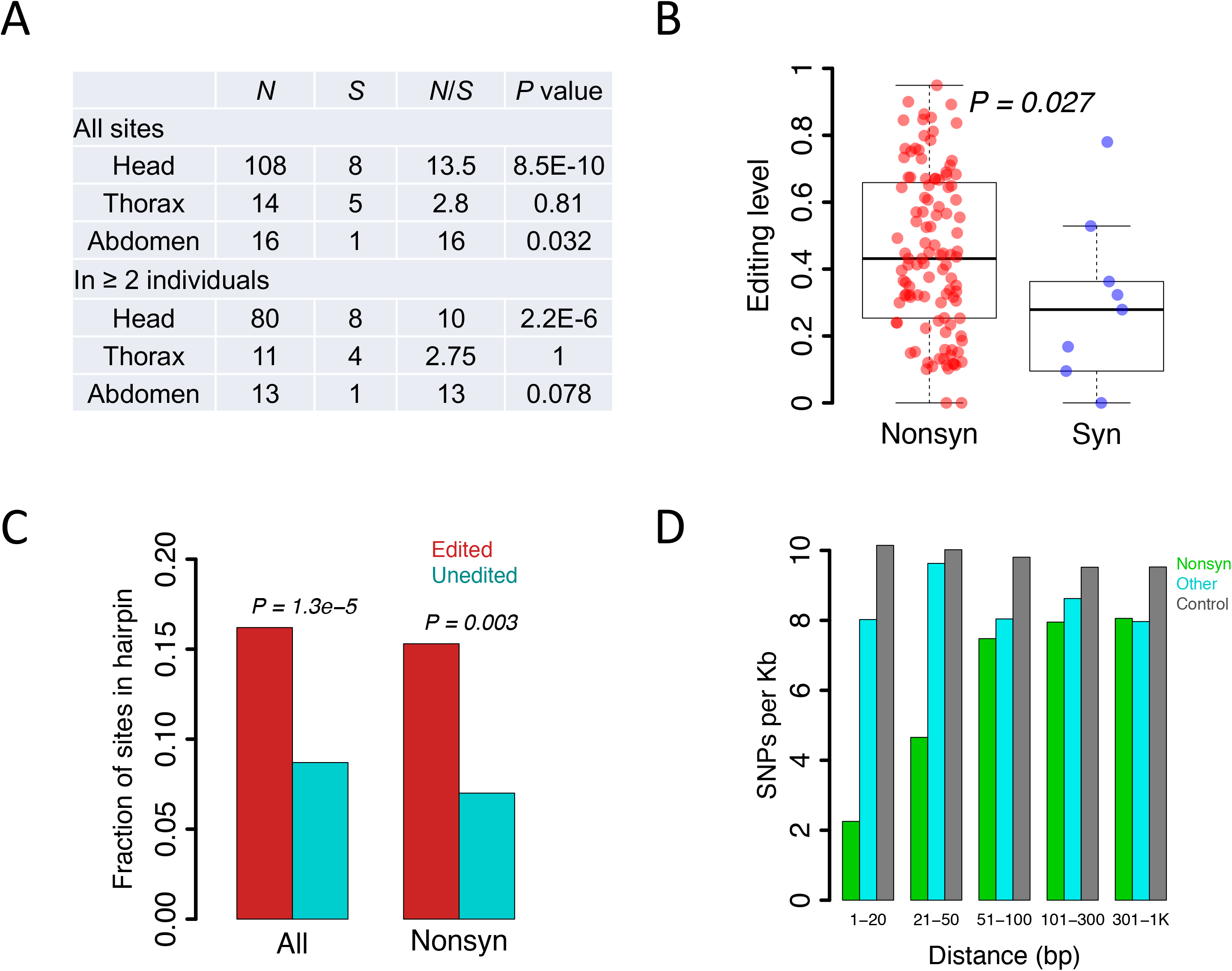
Signals of adaptation of A-to-I editing in honeybee. (A) The observed numbers of nonsynonymous (*N*) and synonymous (*S*) editing sites in each tissue. *P* values are obtained from Fisher’s exact test, comparing the observed *N* and *S* counts to all adenosines in the coding sequence of in *A. mel* (*N*=4,492,737 sites where A-to-G substitution is nonsynonymous, and *S*=1,986,130 for synonymous). “All sites” represents the numbers of sites that appear in at least one individual. Comparisons for sites appearing in at least two individuals are also shown. (B) Editing levels of the nonsynonymous and synonymous sites in head. Data from four drones was pooled to increase the statistical power. *P* values are calculated using Wilcoxon rank sum test. (C) The fraction of sites in predicted stable hairpin structures. The observed editing sites were compared to the unedited adenosines (*P*-values using Fisher’s exact test). (D) The SNP density in the vicinity of nonsynonymous editing sites (green), other editing sites (cyan) or unedited adenosines (grey).

A previous study in cephalopods proposed that the coleoids massively edit their RNAs to diversify the transcriptome at the cost of constraining the evolution of genomic sequences (Liscovitch-Brauer et al. 2017). Maintaining beneficial editing requires genomic conservation of the genomic sequence encoding for the dsRNA structures that allow ADAR to bind and deaminate the adenosine. To test this, we first confirmed that the editing sites in honeybees are enriched in hairpin structures (Figure 2C). Then, we demonstrated that SNPs are depleted in the vicinity of nonsynonymous sites (Figure 2D), further supporting the notion of selective advantage for the nonsynonymous editing events that justifies the tradeoff between transcriptome diversity and genome evolution.

### Evolutionarily conserved and non-conserved editing sites

Editing at four nonsynonymous and one synonymous site is conserved between *Drosophila* and honeybees (Figure 3). Three of these sites are in the *Shab* transcript (*shaker cognate b*, two nonsynonymous and one synonymous) and the other two are is in *qvr* (*quiver*). Editing at these sites is also observed in the bumblebee *Bombus terrestris* (Porath et al. 2019), and editing levels are high in all of these species (Figure 3). These findings suggest a potential functional role for these widely conserved editing sites. However, the majority of the editing sites are not shared between *Drosophila* and honeybees. For some sites the editable adenosine is not conserved in the honeybee genome (Figure S9), and are thus clearly uneditable, while other adenosines are conserved but editing was not detected in RNA-Seq data (Figure S9). Overall, of the 639 *Drosophila* editing sites in CDS for which an orthologous site was identified in honeybees, 349 were conserved as A (55%), and 85, 127, and 78 (13%, 20%, and 12%) were mutated to C, G and T, respectively.

**Figure 3.**
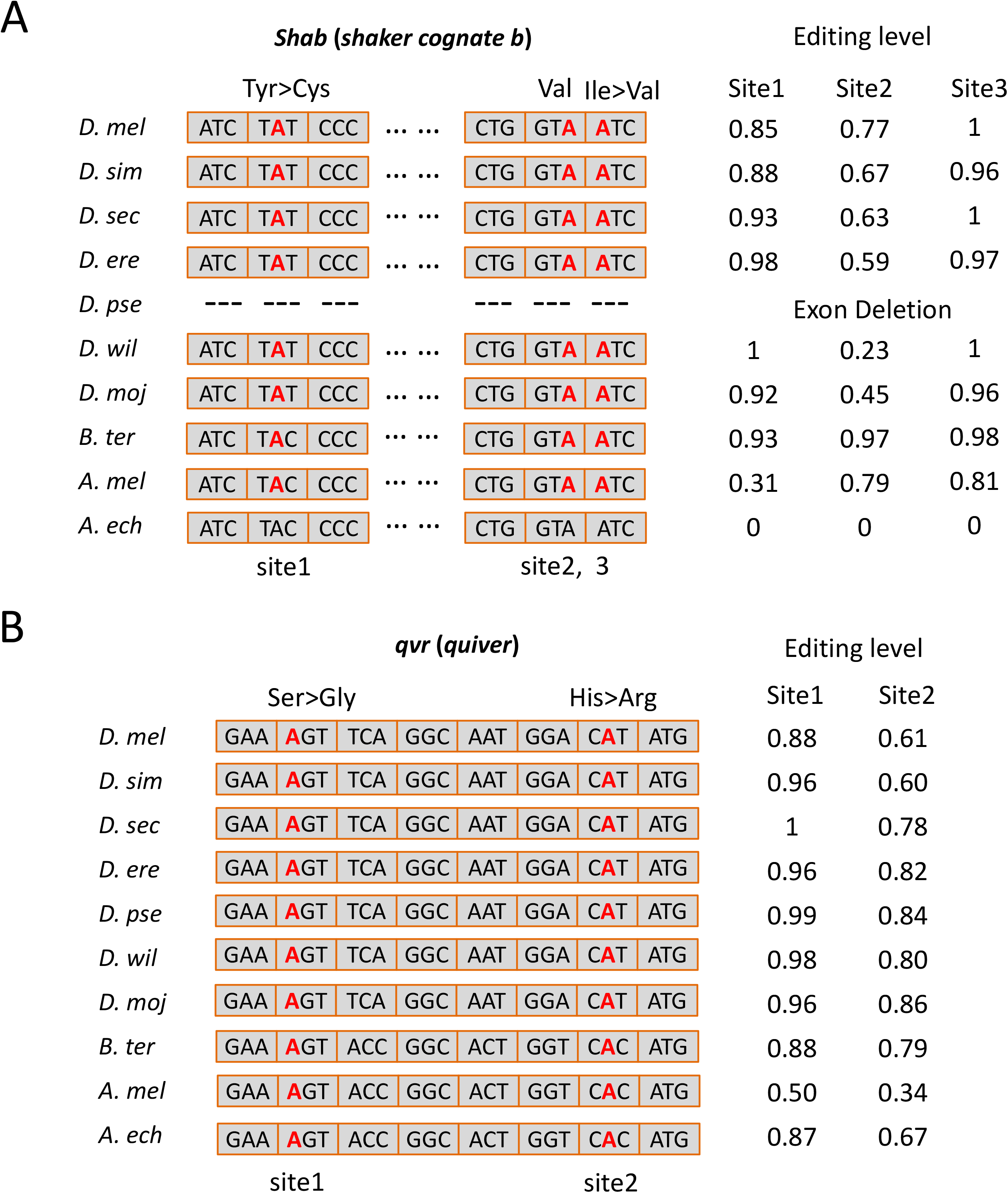
Editing sites conserved between bees and flies. (A) One synonymous and two nonsynonymous editing sites in *Shab* transcripts are edited in heads of two bee and four *Drosophila* species. (B) Two nonsynonymous editing sites in *qvr* transcripts are edited in heads of two bee and four *Drosophila* species. The editing sites are colored red. Editing levels measured in male heads of flies (unpublished), pooled heads of honeybee drones, and brains of bumblebees (Porath et al. 2019) are shown. *D. mel*, *Drosophila melanogaster*; *D. sim*, *Drosophila simulans*; *D. sec*, *Drosophila sechellia*; *D. ere*, *Drosophila erecta; D. pse, Drosophila pseudoobscura; D. wil, Drosophila willistoni; D. moj, Drosophila mojavensis; A. mel*, *Apis mellifera*; *B. ter*, *Bombus terrestris*.

Similarly, most sites are not even conserved across the two bee species. The bumblebee study has reported 219 editing sites in CDS, 164 of which are nonsynonymous (Porath et al. 2019). Of these, we found 9 editing sites in coding regions conserved between honeybee and bumblebee (Table S5).

One of these conserved sites is a Ser>Gly site within the *tipE* (temperature-induced paralytic E; *GB47375*) transcript, editing levels of which correlate with task-performance in bumblebees (nurses versus foragers) (Porath et al. 2019). Editing level at this site in honeybee heads are ~0.7, similar to those observed in bumblebee brains. In the two other tissues this gene is lowly expressed and poorly edited. The orthologous site in *Drosophila* genomes encodes the post-edit Gly codon GGC (Figure S10). Thus, *tipE* recoding seems to be bee-specific, possibly related to social behavior and task performance.

### Sex-dependent and Caste-dependent editing

We further looked for potential sex-dependence and caste-dependent editing, comparing our honeybee drone head data with previously published brain RNA-seq data studying two sub-castes of female workers, foragers and reverted nurses (Herb et al. 2012) (Table S6). First, we compared the pooled editing levels between drones and workers for each well-covered (>10 reads in each pool) editing site separately. Twenty six sites exhibit differential editing (Wilcoxon rank sum test; FDR=0.05) between workers (females) and drones (males), including 4 recoding sites, 2 sites in 3’UTRs, 3 intronic sites, and 17 intergenic sites (Figure 4A and Table S7, Differentially edited sites between drones and workers, Related to Figure 4). In most cases editing in drones is higher, but this may result from looking at sites identified in drone data to begin with. The difference in *Adar* expression was not statistically significant (Figure S11).

**Figure 4.**
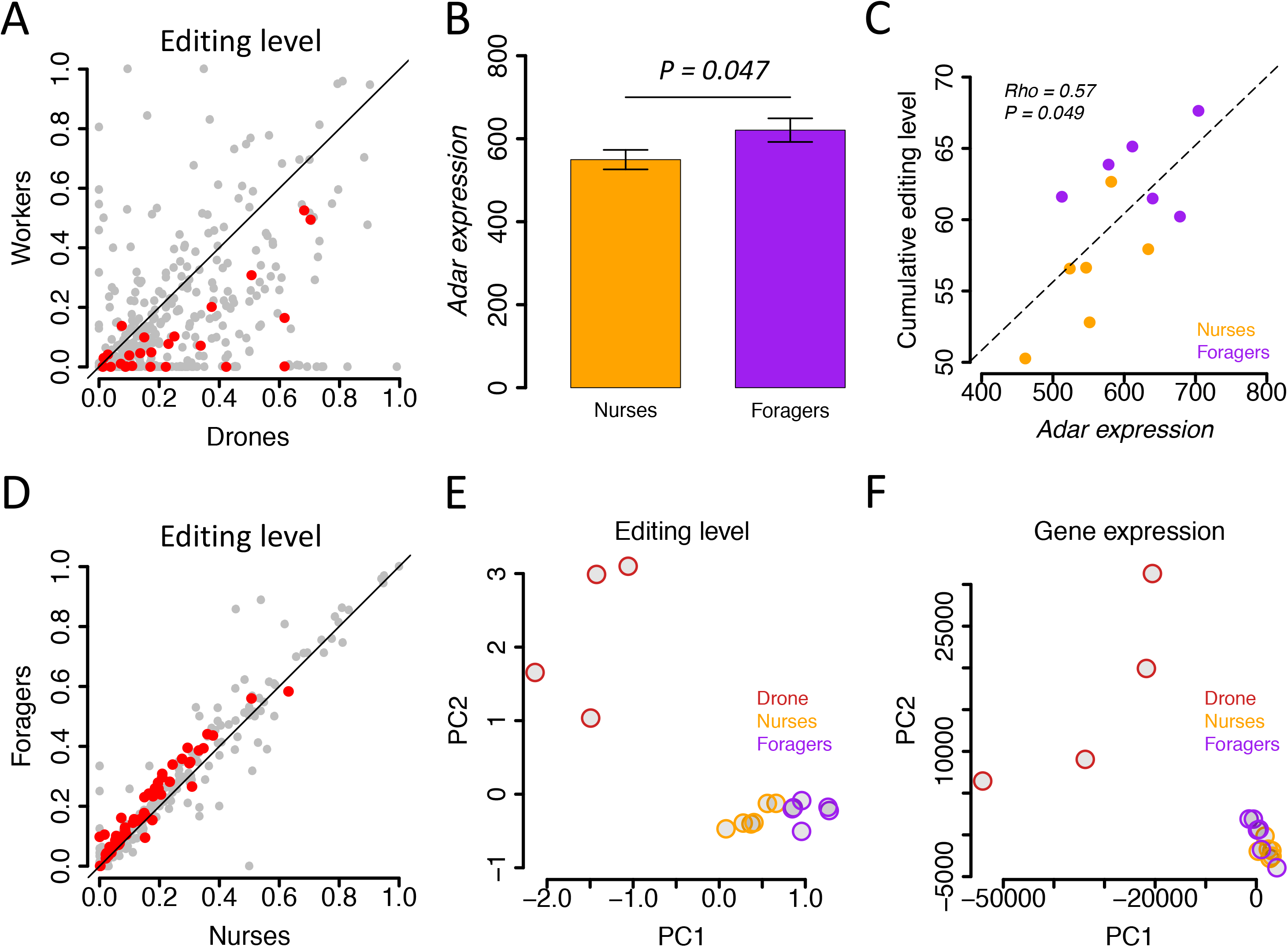
Sex-dependent and caste-dependent editing. (A) Comparison of editing levels between drones (males) and workers (females). Statistically significant sites (FDR < 0.05) are colored red. (B) *Adar* expression level (reads count normalized by DESeq2) in nurses and foragers. Data are presented as mean ± SEM (standard error of mean). *P* value was calculated using Wilcoxon rank sum test. (C) Spearman’s correlation between *Adar* expression and editing index (sum over all G alleles in all sites divided by sum of all the coverages in all sites) for all the nurse (orange) and forager (purple) samples. (D) Comparison of editing levels between two sub-castes of workers: nurses and foragers. Statistically significant sites (FDR < 0.3; see Results) are colored red. (E) PCA analysis of the editing profile shows a different behavior for nurses and foragers, while (F) PCA of the expression profile (genes with RPKM>1) does not lead to a clear separation.

Comparing the pooled editing levels between the two sub-castes (foragers and nurses), one finds 230 out of 407 sites with lower levels in nurses, compared to only 91 exhibiting higher levels in nurses (equal level, mostly zero, was observed in 84 sites), suggesting a globally higher editing activity (at the detected sites) in foragers (*P* = 1.3e-14, proportion test). The surcharge of forager-higher sites is maintained for various coverage and editing-difference cutoffs (Figure S12). Furthermore, *Adar* expression (Figure 4B) and the editing index (Figure 4C) are both higher in foragers, and the editing index correlates with *Adar* expression. However, reliable detection of specific differentially-edited sites with the available sample size (6 foragers and 6 nurses) is challenging, and not even a single site was identified with FDR=0.05 (Wilcoxon rank sum test). We have thus relaxed the statistical test, allowing for FDR=0.3, and found 61 candidate sites, most of which are differentially edited (Figure 4D and Table S8, Differentially edited sites between foragers and nurses, Related to Figure 4). These include 10 nonsynonymous, 16 3’UTR, and 35 intergenic sites. Consistently, 55 of these sites exhibit higher levels of editing in foragers. Finally, PCA analysis of the editing profile across sites results in distinct clusters of drones, nurses and foragers, a classification that is not achieved by the expression profile (Figure 4E and 4F). Taken together, these results show a global increase in editing in foragers compared to nurses.

### Convergent adaptation of A-to-I editing?

One of the recoding sites conserved between the two bee species resides within *Adar* transcript (Table S5 and Figure 5A). In bumblebee, the recoding level of the conserved site, I482M (ATA to ATG), positively correlates with the global editing activity (Porath et al. 2019), suggesting a possible auto-regulation mechanism. Interestingly, *Drosophila Adar* is also auto-edited at a different position, where a Ser (AGT) to Gly (GGT) substitution leads to a less active ADAR protein, resulting in a negative feedback loop of editing activity (Palladino et al. 2000; Savva et al. 2012a). Intriguingly, fly-edited Ser amino-acid is conserved in honeybee and bumblebee, but a different codon (TCA, uneditable at the first codon position) is used. Similarly, the bee-edited Ile is conserved in flies, but the edited adenosines (at the third position of ATA) is synonymously mutated to T, and the editable ATA bee codon is substituted by ATT codon in flies, which could not be edited at the third position (Figure 5A). Thus, while auto-editing S430G in flies is abolished in bees due to an uneditable Ser codon, and the auto-editing I482M in bees is abolished in flies due to an uneditable Ile codon. However, the mechanism of auto-editing, possibly used for global ADAR regulation, is shared by the two lineages, albeit at different positions and possibly with different effects on the protein. This might hint at the possibility of convergent evolution of ADAR auto-regulation strategy.

**Figure 5.**
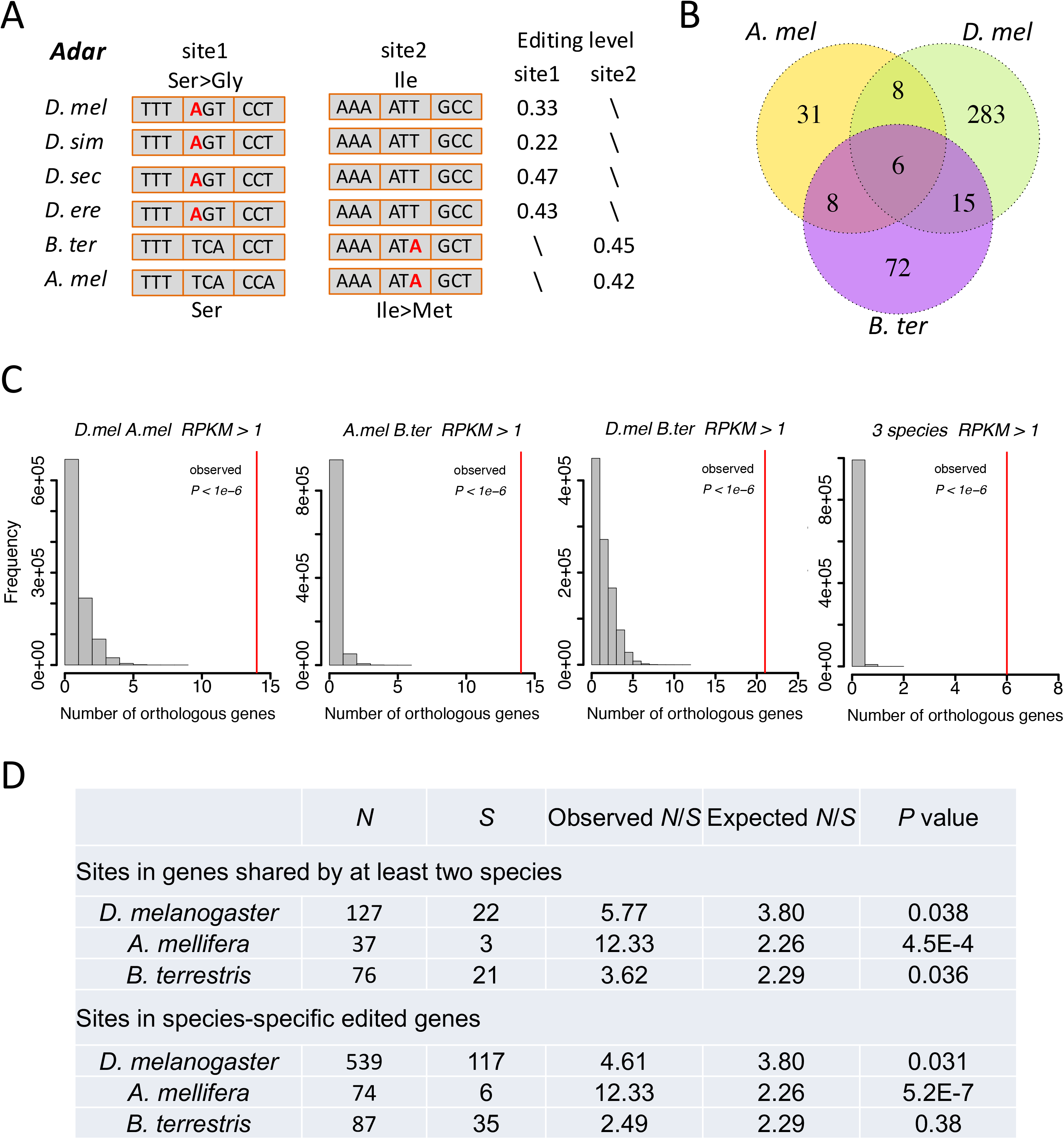
Convergent evolution of editing. (A) *Adar* transcripts are auto-edited in *Drosophila* and bees. The Ser>Gly site is highly conserved across *Drosophila* species. The *Drosophila*-editable Ser codon AGT is changed to an uneditable Ser codon TCA in bees. On the other hand, the Ile>Met auto-editing site is conserved between honeybee and bumblebee. The bee-editable Ile codon ATA (edited at the third position) appears as an uneditable Ile codon ATT in flies. Editing sites are colored red. The editing levels from male heads of flies (unpublished), pooled heads of honeybee drones and brains of bumblebee (Porath et al. 2019) are shown. *D. mel*, *Drosophila melanogaster*; *D. sim*, *Drosophila simulans*; *D. sec*, *Drosophila sechellia*; *D. ere*, *Drosophila erecta*; *A. mel*, *Apis mellifera*; *B. ter*, *Bombus terrestris*. (B) Venn diagram demonstrating the overlap between orthologous genes with editing sites in coding regions. (C) The observed number of orthologous genes with editing in coding regions, compared to the expected distribution. Genes with RPKM > 1 in both species were chosen to calculate the expected numbers (results are essentially the same for other expression cutoffs, Figure S13). *P*-values calculated by randomization test. (D) The nonsynonymous to synonymous (*N*/*S*) ratios are significantly higher than expected, for both shared and species-specific sites (except for species-specific sites in *B. terrestris*). Here the shared and species-specific sites refer to the sites in shared genes and species-specific genes as defined in the above Venn diagram in (B). The expected ratio is evaluated by calculating the effect of putative editing on all adenosines in the coding region. *P*-values by Fisher exact test.

Following this example, we wondered whether there are more genes for which editing is observed in both honeybees and *Drosophila*, even if the exact location of the editing site is not conserved. Excluding conserved sites, there are 53 genes exhibiting CDS editing in honeybee heads, 101 genes exhibiting CDS editing in bumblebee heads, and 312 genes edited in CDS in male brains of *D. melanogaster*. Of these, 21 genes exhibit editing in both bee species, 14 are edited in honeybee and *Drosophila,* 14 are edited in bumblebee and *Drosophila*, and six genes are found to be edited in all three species (Figure 5B and Table S9). These numbers are significantly higher than expected by random sampling, (p<1e-6, randomization test), even if one accounts for the expression profile (Transparent Methods, Figure 5C and Figure S13). Table S9). We have repeated the analysis using stricter criteria to characterize sites, considering as edited only sites with an observed editing level >10% (all samples pooled), and a binomial *P* < 0.05 (excluding the null hypothesis of no editing), and considering a site to be not-edited if the binomial *P* < 0.05 for the null hypothesis that the site is edited at 10%. Reassuringly, the results are robust to this change of definitions (Figure S14 and S15). These results suggest an interesting convergent evolution of A-to-I editing. Recoding sites are rarely conserved across clades, but the same genes in different lineages tend to acquire a recoding event, apparently independently, indicating a functional importance for recoding of this gene.

## DISCUSSION

Despite over fifteen years of developments, reliable detection of CDS editing remains a challenge. Systematic searches often lead to high false-positive rates (reflected by the fraction of non-AG mismatches observed), especially for mammalian transcriptomes where the scope of recoding is rather low. Important exception are *Drosophila* and cephalopods, where the recoding signal is much more pronounced and easier to detect. Applying a novel strict alignment approach, we were able to achieve here a high accuracy in CDS editing detection (84.3% in heads) despite the overall modest scope of recoding sites. Another similarity between honeybees *Drosophila* and cephalopods is related to the important question regarding the extent to which recoding is adaptive. This is often estimated by the *N/S* ratio, where *Drosophila* and cephalopods exhibit a pattern markedly different from the one seen in other species studied, (including human). Whereas in most species the *N/S* ratio observed is lower or similar to the one expected under neutrality, it is significantly higher in *Drosophila* and cephalopods. Again, we find here that honeybees show a strikingly high *N/S* ratio, similar to the above two clades. We suggest that the two last points are interconnected. The apparently low *N/S* ratio observed in many species (human included) may reflect the low accuracy of the lists of CDS editing sites and not the (lack of) adaptive potential for recoding in these species.

The importance of post-transcriptional and post-translational mechanisms in generating the proteomic complexity of higher organisms has been emphasized in recent decades. These epigenetic mechanisms allow for diversification of the proteome and functional heterogeneity across tissues, developmental stages, brain regions or even among individual cells.

Recoding by A-to-I RNA editing is an epigenetic mechanism capable of diversifying the proteome, creating a range of proteins from a single genomically encoded gene in a temporally-regulated, tissue-specific, condition-dependent way, and providing the organism with a new means for acclimation and adaptation. Indeed, several studies have demonstrated how recoding levels at specific sites do change as a function of the organism’s condition (Garrett and Rosenthal 2012; Robinson et al. 2016; Gallo et al. 2017; Porath 2017; Terajima et al. 2017; Yablonovitch et al. 2017). Importantly, many studies have demonstrated altered editing of individual recoding targets in various disease states (Gallo et al. 2017). However, these interesting examples notwithstanding, the extent to which the recoding phenomenon is actually used as a means for proteome diversification is still under debate. In fact, several recent studies have raised the possibility that nonsynonymous editing may be used to compensate for otherwise deleterious G-to-A mutations, rather than allowing for the two alleles (A and G) to co-exist (Jiang and Zhang 2019; Mai and Chuang 2019; Popitsch et al. 2020). Thus, a high *N*/*S* ratio is not sufficient to prove that editing serves for adaptation through proteome diversification.

The behavioral, physiological and morphological plasticity exhibited by social insects suggests an important role for epigenetic mechanisms. Differential RNA editing was previously shown for different castes of the highly social leaf-cutting ant *Acromyrmex echinatior* (Li et al. 2014), as well as for bumblebee workers (Porath et al. 2019). Our results expand these finding to another species, further supporting the notion of RNA editing as a source of proteomic complexity, that may be recruited to provide phenotypic variation among genetically identical individuals.

ADAR enzymes, and the recoding phenomenon are widely conserved across metazoan. However, the repertoire of recoding sites varies considerably and seems to have developed almost independently in different clades. Here we point out that the same gene targets are being edited in distant species, even if the exact locations of the editing sites within the transcript vary. A particularly interesting example is that of nonsynonymous auto-editing of *Adar*, shown to affect ADAR activity in bumblebees, flies and even mammals (Rueter et al. 1999; Savva et al. 2012a; Porath et al. 2019), possibly serving as global editing autoregulation. It thus seems possible that the selective advantage in having multiple versions of the protein product is shared by the target genes, while there is more than one specific way to achieve this diversity through recoding. Clades developed along different evolutionary routes have converged on different implementations (in terms of the specific editing site) of the same diversifying solution. These ideas should be further tested in the future, using larger datasets and across additional species and clades. Moreover, as biochemical and functional understanding of the impact of the different edits are gained, it will be possible to look into a possible common effect for the different edits of the common targets.

### Limitations of the Study

The editomes of the bee species analyzed here are based on a limited number of samples. They probably represent a subset of the actual repertoire of editing sites. The results should be re-tested when more data is available. In particular, the honeybee editome was built using drone samples. Thus, some individual differential editing between the nurses and foragers sub-castes may have missed worker-specific sites. However, the global increased editing in foragers, supported by a higher Adar expression, is probably robust.

### Resource Availability

#### Lead Contact

Further information and requests for resources should be directly and will be fulfilled by the Lead Contact, Jian Lu.

#### Materials Availability

This study did not generate new unique materials.

#### Data and Code Availability

All deep-sequencing data generated in this study were deposited in the China National Genomics Data Center Genome Sequence Archive (GSA) under accession number CRA002262. Other deep-sequencing data analyzed were downloaded from SRA as follows. Brains of honeybee workers: accession numbers SRR445999 to SRR446004 (reverted nurse) and SRR446005 to SRR446010 (forager) (Herb et al. 2012). Bumblebee: SRP166322 (Porath et al. 2019). *D. mel* head: SRP067542 (Zhang et al. 2018). *D. sim* head: SRP074828 (Duan et al. 2017). *D. pse*: DRR055250 and DRR055251 (Nozawa et al. 2016). *D. wil*: SRR341127 and SRR341129 (Meisel et al. 2012). *D. moj*: SRR037508-SRR037518 (generated by the *Drosophila* modENCODE project). *D. sec*, and *D. ere* data are unpublished data generated in our own lab. The reference genomes versions used for *Drosophila* species are Dmel_r6.04 (http://flybase.org/) for *D. mel*, and versions droSim1, droSec1, droEre1, dp4, droWil1, and droMoj3 downloaded from UCSC Genome Browser (http://genome.ucsc.edu/) for the other fly species. For the *Bombus terrestris*, we used version Bter_1.0.

## METHODS

All methods can be found in the accompanying Transparent Methods supplemental file.

## ACKNOWLEDGMENTS

We thank the National Center for Protein Sciences at Peking University for technical assistance. Part of the analysis was performed on the High Performance Computing Platform of the Center for Life Science at Peking University. This work was supported by the joint NSFC-ISF program (NSFC grant number 3201101147 to J.L. and ISF grant number 3371/20 to E.E.), as well as grants from the Ministry of Science and Technology of the People’s Republic of China (2016YFA0500800) and National Natural Science Foundation of China (91731301) to J.L. and support from the Israel Science Foundation (1945/18) to E.E.

## AUTHOR CONTRIBUTIONS

J.L. designed the research; J.X.H. contributed the honeybee materials; S.Q.D. performed the research; Y.G.D., H.T.P, E.E. and J.L. performed the bioinformatics analyses; E.E. and J.L. wrote the paper.

## DECLARATION OF INTERESTS

The authors declare no competing interests.

Nonsynonymous editing sites in honeybees were under positive selection

Differential editing may contribute to the phenotypic diversity between sub-castes

Target genes acquire editing in different clades, suggesting convergent evolution

## Notes

https://bigd.big.ac.cn/gsa/browse/CRA002262

### Competing Interest Statement

The authors have declared no competing interest.

### Summary of Updates

This new version has been extensively updated from the previous version. This study has been published by iScience at https://doi.org/10.1016/j.isci.2020.101983

## REFERENCES

Alon S, Garrett SC, Levanon EY, Olson S, Graveley BR, Rosenthal JJ, Eisenberg E. 2015. The majority of transcripts in the squid nervous system are extensively recoded by A-to-I RNA editing. Elife 4.

Alon S, Mor E, Vigneault F, Church GM, Locatelli F, Galeano F, Gallo A, Shomron N, Eisenberg E. 2012. Systematic identification of edited microRNAs in the human brain. Genome Res 22: 1533–1540.

Athanasiadis A, Rich A, Maas S. 2004. Widespread A-to-I RNA editing of Alu-containing mRNAs in the human transcriptome. PLoS Biol 2: e391.

Basilio C, Wahba AJ, Lengyel P, Speyer JF, Ochoa S. 1962. Synthetic polynucleotides and the amino acid code. V. Proc Natl Acad Sci U S A 48: 613–616.

Bass BL. 2002. RNA editing by adenosine deaminases that act on RNA. Annu Rev Biochem 71: 817–846.

Blow M, Futreal PA, Wooster R, Stratton MR. 2004. A survey of RNA editing in human brain. Genome Research 14: 2379–2387.

Borchert GM, Gilmore BL, Spengler RM, Xing Y, Lanier W, Bhattacharya D, Davidson BL. 2009. Adenosine deamination in human transcripts generates novel microRNA binding sites. Human Molecular Genetics 18: 4801–4807.

Buchumenski I, Bartok O, Ashwal-Fluss R, Pandey V, Porath HT, Levanon EY, Kadener S. 2017. Dynamic hyper-editing underlies temperature adaptation in Drosophila. PLOS Genetics 13: e1006931.

Chen J-Y, Peng Z, Zhang R, Yang X-Z, Tan BC-M, Fang H, Liu C-J, Shi M, Ye Z-Q, Zhang YE et al. 2014. RNA Editome in Rhesus Macaque Shaped by Purifying Selection. PLoS Genet 10: e1004274.

Danecek P, Nellaker C, McIntyre RE, Buendia-Buendia JE, Bumpstead S, Ponting CP, Flint J, Durbin R, Keane TM, Adams DJ. 2012. High levels of RNA-editing site conservation amongst 15 laboratory mouse strains. Genome Biol 13: 26.

Duan Y, Dou S, Luo S, Zhang H, Lu J. 2017. Adaptation of A-to-I RNA editing in Drosophila. PLoS Genet 13: e1006648.

Eisenberg E, Levanon EY. 2018. A-to-I RNA editing — immune protector and transcriptome diversifier. Nature Reviews Genetics 19: 473–490.

Flomen R, Knight J, Sham P, Kerwin R, Makoff A. 2004. Evidence that RNA editing modulates splice site selection in the 5-HT2C receptor gene. Nucleic Acids Research 32: 2113–2122.

Gallo A, Vukic D, Michalik D, O’Connell MA, Keegan LP. 2017. ADAR RNA editing in human disease; more to it than meets the I. Hum Genet 136: 1265–1278.

Garrett S, Rosenthal JJ. 2012. RNA editing underlies temperature adaptation in K+ channels from polar octopuses. Sci 335: 848–851.

Goldstein B, Agranat-Tamir L, Light D, Ben-Naim Zgayer O, Fishman A, Lamm AT. 2017. A-to-I RNA editing promotes developmental stage–specific gene and lncRNA expression. Genome Research 27: 462–470.

Gommans WM, Mullen SP, Maas S. 2009. RNA editing: a driving force for adaptive evolution? BioEssays 31: 1137–1145.

Graveley BR, Brooks AN, Carlson JW, Duff MO, Landolin JM, Yang L, Artieri CG, van Baren MJ, Boley N, Booth BW et al. 2011. The developmental transcriptome of Drosophila melanogaster. Nature 471: 473–479.

Herb BR, Wolschin F, Hansen KD, Aryee MJ, Langmead B, Irizarry R, Amdam GV, Feinberg AP. 2012. Reversible switching between epigenetic states in honeybee behavioral subcastes. Nat Neurosci 15: 1371–1373.

Jiang D, Zhang J. 2019. The preponderance of nonsynonymous A-to-I RNA editing in coleoids is nonadaptive. Nat Commun 10: 5411.

Jin Y, Tian N, Cao J, Liang J, Yang Z, Lv J. 2007. RNA editing and alternative splicing of the insect nAChR subunit alpha6 transcript: evolutionary conservation, divergence and regulation. BMC Evolutionary Biology 7: 1–12.

Keegan LP, Gallo A, O’Connell MA. 2001. The many roles of an RNA editor. Nat Rev Genet 2: 869–878.

Keegan LP, McGurk L, Palavicini JP, Brindle J, Paro S, Li X, Rosenthal JJ, O’Connell MA. 2011. Functional conservation in human and Drosophila of Metazoan ADAR2 involved in RNA editing: loss of ADAR1 in insects. Nucleic Acids Res 39: 7249–7262.

Kim DD, Kim TT, Walsh T, Kobayashi Y, Matise TC, Buyske S, Gabriel A. 2004. Widespread RNA editing of embedded alu elements in the human transcriptome. Genome Res 14: 1719–1725.

Klironomos FD, Berg J, Collins S. 2013. How epigenetic mutations can affect genetic evolution: Model and mechanism. BioEssays 35: 571–578.

Lev-Maor G, Sorek R, Levanon EY, Paz N, Eisenberg E, Ast G. 2007. RNA-editing-mediated exon evolution. Genome biology 8: R29.

Levanon EY, Eisenberg E, Yelin R, Nemzer S, Hallegger M, Shemesh R, Fligelman ZY, Shoshan A, Pollock SR, Sztybel D et al. 2004. Systematic identification of abundant A-to-I editing sites in the human transcriptome. Nat Biotech 22: 1001–1005.

Li Q, Wang Z, Lian J, Schiott M, Jin L, Zhang P, Zhang Y, Nygaard S, Peng Z, Zhou Y et al. 2014. Caste-specific RNA editomes in the leaf-cutting ant Acromyrmex echinatior. Nat Commun 5: 4943.

Liang H, Landweber LF. 2007. Hypothesis: RNA editing of microRNA target sites in humans? RNA 13: 463–467.

Licht K, Hartl M, Amman F, Anrather D, Janisiw MP, Jantsch MF. 2019. Inosine induces context-dependent recoding and translational stalling. Nucleic Acids Res 47: 3–14.

Liscovitch-Brauer N, Alon S, Porath HT, Elstein B, Unger R, Ziv T, Admon A, Levanon EY, Rosenthal JJC, Eisenberg E. 2017. Trade-off between Transcriptome Plasticity and Genome Evolution in Cephalopods. Cell 169: 191–202 e111.

Mai TL, Chuang TJ. 2019. A-to-I RNA editing contributes to the persistence of predicted damaging mutations in populations. Genome Res 29: 1766–1776.

Mazloomian A, Meyer IM. 2015. Genome-wide identification and characterization of tissue-specific RNA editing events in D. melanogaster and their potential role in regulating alternative splicing. RNA Biology 12: 1391–1401.

Meisel RP, Malone JH, Clark AG. 2012. Disentangling the relationship between sex-biased gene expression and X-linkage. Genome Res 22: 1255–1265.

Morse DP, Bass BL. 1999. Long RNA hairpins that contain inosine are present in Caenorhabditis elegans poly(A)+ RNA. Proc Natl Acad Sci U S A 96: 6048–6053.

Neeman Y, Levanon EY, Jantsch MF, Eisenberg E. 2006. RNA editing level in the mouse is determined by the genomic repeat repertoire. RNA 12: 1802–1809.

Nishikura K. 2010. Functions and regulation of RNA editing by ADAR deaminases. Annu Rev Biochem 79: 321–349.

Nishikura K. 2016. A-to-I editing of coding and non-coding RNAs by ADARs. Nat Rev Mol Cell Biol 17: 83–96.

Nozawa M, Onizuka K, Fujimi M, Ikeo K, Gojobori T. 2016. Accelerated pseudogenization on the neo-X chromosome in Drosophila miranda. Nat Commun 7: 13659.

Page RE Jr., Rueppell O, Amdam GV. 2012. Genetics of reproduction and regulation of honeybee (Apis mellifera L.) social behavior. Annu Rev Genet 46: 97–119.

Palladino MJ, Keegan LP, O’Connell MA, Reenan RA. 2000. dADAR, a Drosophila double-stranded RNA-specific adenosine deaminase is highly developmentally regulated and is itself a target for RNA editing. RNA 6: 1004–1018.

Picardi E, D’Erchia AM, Lo Giudice C, Pesole G. 2017. REDIportal: a comprehensive database of A-to-I RNA editing events in humans. Nucleic Acids Res 45: D750–D757.

Popitsch N, Huber CD, Buchumenski I, Eisenberg E, Jantsch M, von Haeseler A, Gallach M. 2020. A-to-I RNA Editing Uncovers Hidden Signals of Adaptive Genome Evolution in Animals. Genome Biol Evol 12: 345–357.

Porath HT. 2017. A-To-I RNA editing in the earliest-diverging eumetazoan phyla. Mol Biol Evol 34.

Porath HT, Hazan E, Shpigler H, Cohen M, Band M, Ben-Shahar Y, Levanon EY, Eisenberg E, Bloch G. 2019. RNA editing is abundant and correlates with task performance in a social bumblebee. Nature Communications 10.

Porath HT, Knisbacher BA, Eisenberg E, Levanon EY. 2017a. Massive A-to-I RNA editing is common across the Metazoa and correlates with dsRNA abundance. Genome Biol 18: 185.

Porath HT, Schaffer AA, Kaniewska P, Alon S, Eisenberg E, Rosenthal J, Levanon EY, Levy O. 2017b. A-to-I RNA Editing in the Earliest-Diverging Eumetazoan Phyla. Molecular Biology and Evolution 34: 1890–1901.

Ramaswami G, Li JB. 2016. Identification of human RNA editing sites: A historical perspective. Methods 107: 42–47.

Robinson JE, Paluch J, Dickman DK, Joiner WJ. 2016. ADAR-mediated RNA editing suppresses sleep by acting as a brake on glutamatergic synaptic plasticity. Nature Communications 7.

Rodriguez J, Menet JS, Rosbash M. 2012. Nascent-seq indicates widespread cotranscriptional RNA editing in Drosophila. Mol Cell 47: 27–37.

Rosenthal JJC. 2015. The emerging role of RNA editing in plasticity. The Journal of Experimental Biology 218: 1812.

Rueter SM, Dawson TR, Emeson RB. 1999. Regulation of alternative splicing by RNA editing. Nature 399: 75–80.

Savva YA, Jepson JE, Sahin A, Sugden AU, Dorsky JS, Alpert L, Lawrence C, Reenan RA. 2012a. Auto-regulatory RNA editing fine-tunes mRNA re-coding and complex behaviour in Drosophila. Nat Commun 3: 790.

Savva YA, Rieder LE, Reenan RA. 2012b. The ADAR protein family. Genome Biol 13.

St Laurent G, Tackett MR, Nechkin S, Shtokalo D, Antonets D, Savva YA, Maloney R, Kapranov P, Lawrence CE, Reenan RA. 2013. Genome-wide analysis of A-to-I RNA editing by single-molecule sequencing in Drosophila. Nat Struct Mol Biol 20: 1333–1339.

Terajima H, Yoshitane H, Ozaki H, Suzuki Y, Shimba S, Kuroda S, Iwasaki W, Fukada Y. 2017. ADARB1 catalyzes circadian A-to-I editing and regulates RNA rhythm. Nat Genet 49: 146–151.

Xu G, Zhang J. 2014. Human coding RNA editing is generally nonadaptive. Proc Natl Acad Sci U S A 111: 3769–3774.

Yablonovitch AL, Fu J, Li K, Mahato S, Kang L, Rashkovetsky E, Korol AB, Tang H, Michalak P, Zelhof AC et al. 2017. Regulation of gene expression and RNA editing in Drosophila adapting to divergent microclimates. Nature Communications 8: 1570.

Yan H, Simola DF, Bonasio R, Liebig J, Berger SL, Reinberg D. 2014. Eusocial insects as emerging models for behavioural epigenetics. Nat Rev Genet 15: 677–688.

Yang X-Z, Chen J-Y, Liu C-J, Peng J, Wee YR, Han X, Wang C, Zhong X, Shen QS, Liu H et al. 2015. Selectively Constrained RNA Editing Regulation Crosstalks with piRNA Biogenesis in Primates. Molecular Biology and Evolution 32: 3143–3157.

Yu Y, Zhou H, Kong Y, Pan B, Chen L, Wang H, Hao P, Li X. 2016. The Landscape of A-to-I RNA Editome Is Shaped by Both Positive and Purifying Selection. PLoS Genet 12: e1006191.

Zhang H, Dou S, He F, Luo J, Wei L, Lu J. 2018. Genome-wide maps of ribosomal occupancy provide insights into adaptive evolution and regulatory roles of uORFs during Drosophila development. PLoS Biol 16: e2003903.

Zhang R, Deng P, Jacobson D, Li JB. 2017. Evolutionary analysis reveals regulatory and functional landscape of coding and non-coding RNA editing. PLoS genetics 13: e1006563.

Zhao HQ, Zhang P, Gao H, He X, Dou Y, Huang AY, Liu XM, Ye AY, Dong MQ, Wei L. 2015. Profiling the RNA editomes of wild-type C. elegans and ADAR mutants. Genome Res 25: 66–75.

