## Supplementary material for "A-to-I RNA editing in honeybees shows signals of adaptation and convergent evolution": Methods, and Supplementary table and figures

**Supplemental Figures**

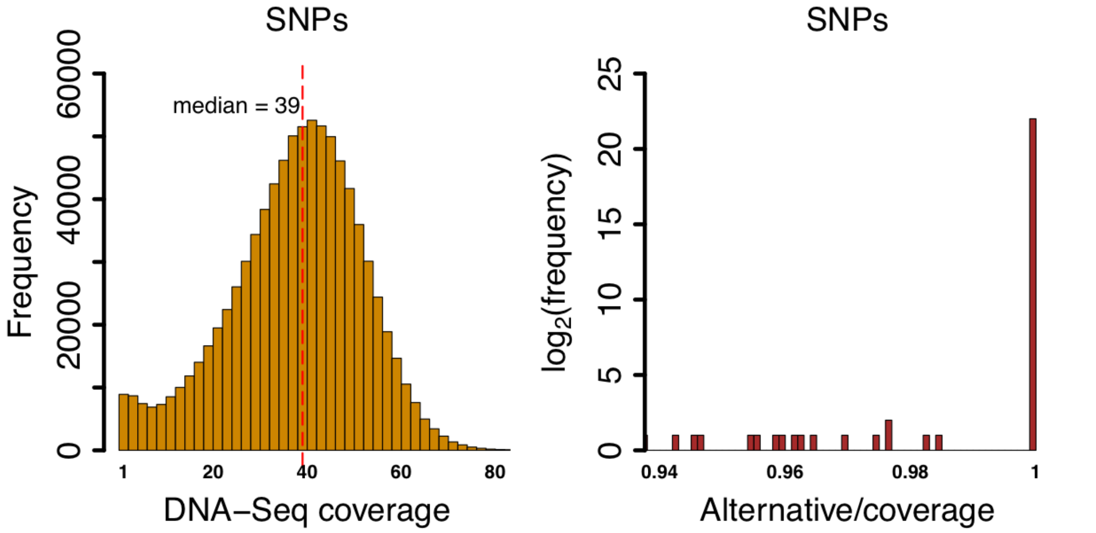


**Figure S1. Per-individual DNA-Seq coverage at detected SNPs sites (left) and the variation level (alternative reads count / coverage) of these SNPs (right), Related to Figure1.**

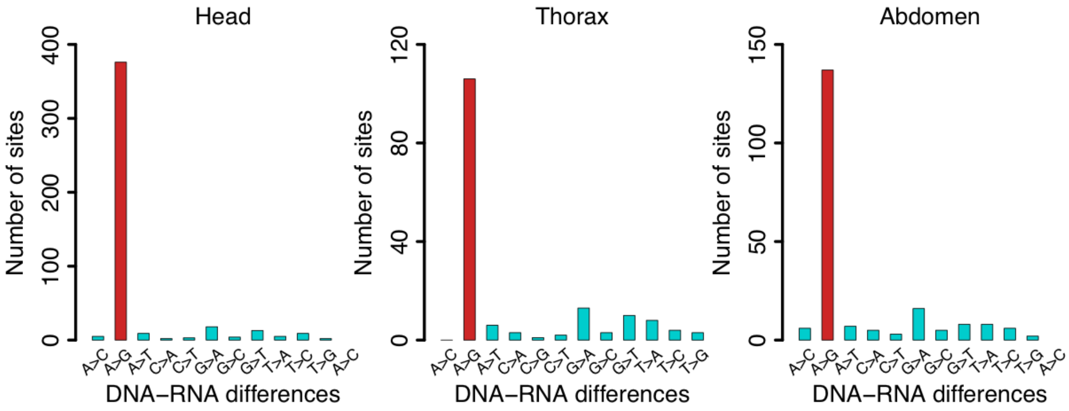


**Figure S2. Distribution of DNA-RNA mismatch types for sites detected in heads, thoraxes, or abdomens of honeybees, Related to Figure 1.**

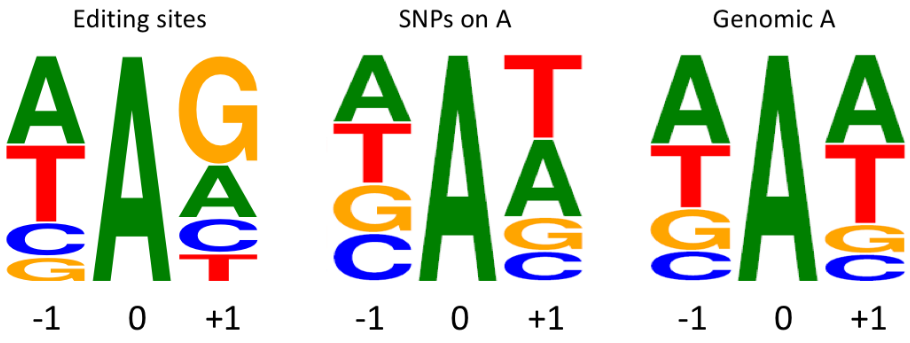


**Figure S3. Local sequence motif around editing sites (left), SNPs (middle) or unedited adenosines in genome (right), Related to Figure 1.**

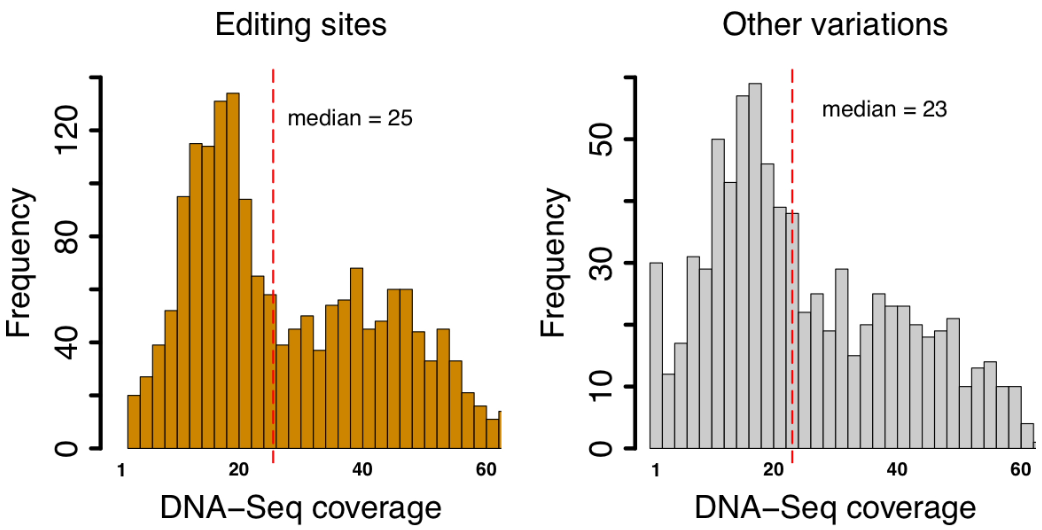


**Figure S4. Histogram of per-individual DNA-Seq coverage at candidate RNA editing sites and other variations in each sample, Related to Figure 1.**

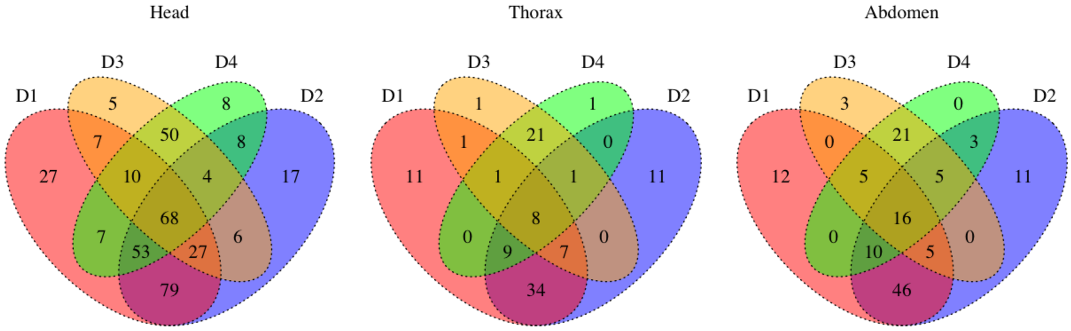


**Figure S5. Venn diagram demonstrating the overlaps of editing sites in four honeybee drones, Related to Figure 1.**

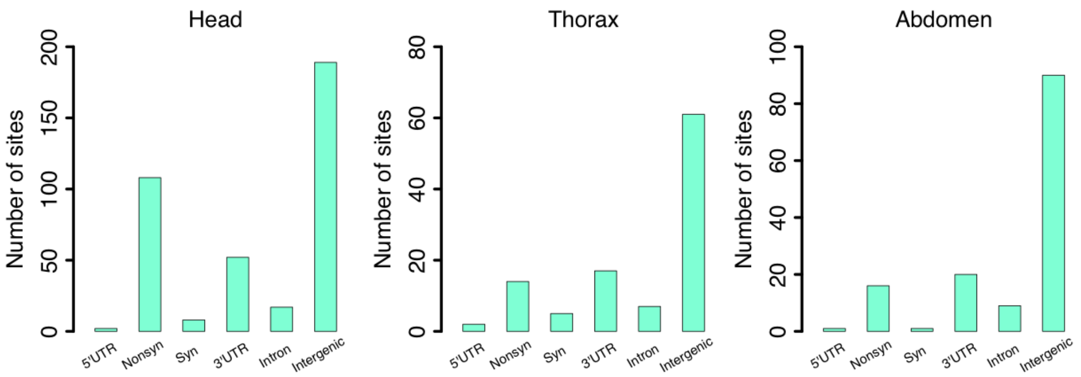


**Figure S6. Functional annotation of editing sites detected in heads, thoraxes, or abdomens of honeybees, Related to Figure 1.**

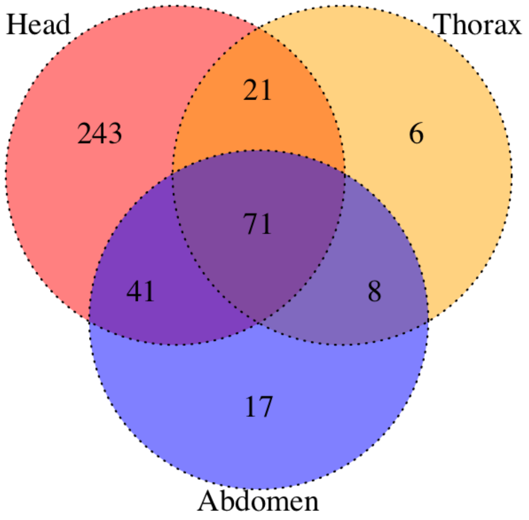


**Figure S7. Venn diagram demonstrating the overlaps of editing sites in three honeybee tissues, Related to Figure 1.**

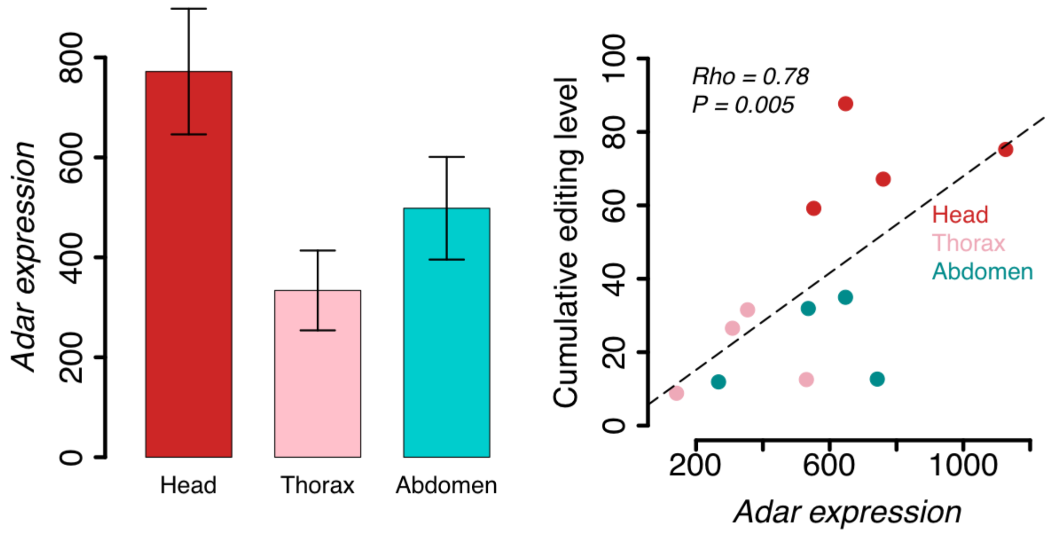


**Figure S8.** ***Adar*** **expression is highest in heads (left) and is positively correlated (Spearman's correlation) to cumulative editing levels in each sample (right), Related to Figure 1.** The cumulative editing level is the sum of the editing level over all sites. Error bars represent the standard error of mean. *Adar* expression values were calculated by the DESeq2 software.

***NaCP60E*** CDS (chr2R:24,914,681-24,914,878)

| | |
|---|---|
| *D. mel* | **… TTCTCGCAACTGTCCGATTT TATTGCC … ATACAC … ACCGA TAACTTCAAACAGTTGCAGGAA …** |
| *B. ter* | **… TTCTCGCAATTAAGCGATTTCATAGCA … ATTCAT … TCTGAAGATTTCCGAAAGCTACAGGAA …** |
| *A. mel* | **… TTTTCGCAATTAAGCGATTTCATAGCA … ATTCAT … TCTGAAGATTTTCGAAAGCTGCAGGAA …** |

***eag*** CDS (chr X:14,999,867-14,999,938)

| | |
|---|---|
| *D. mel* | **… TATTGTCCGAAAGATATGAAGGCTGACATATG TGTTCA TCTAAA TCGC AAAGTATTTAACGAGCAT CCGGCA …** |
| *B. ter* | **… TATTGCCCCAAGGAT ATGAAGGCGGATATCTGCGTTCA TCTGAACAGGAAGGTCTTCAACGAACACCC TGCA …** |
| *A. mel* | **… TACTGCCCTAAAGACATGAAGGCGGACATTTGCGTTCACCTGAACAGGAAGGTGTTCAACGAGCATCC TGCG …** |

**Figure S9. Non-conserved editing sites in gene *NaCP60E* and *eag*, Related to Figure 3.**

*Drosophila*-specific editing sites are colored in red**.** Many of these sites are genomically encoded as G in the honeybee genome.

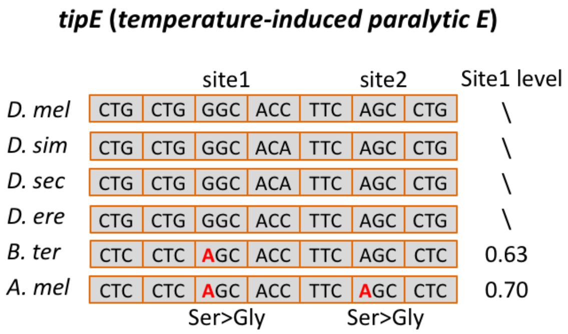


**Figure S10. Bee-specific editing sites in *tipE*, Related to Figure 3.** Two *tipE* recoding sites are observed in honeybee (orthologous gene *GB47375*), one of which is also seen in bumblebee (orthologous gene *XM_003393369*) (Porath et al. 2019), colored in red. Editing is not seen at this site in *Drosophila*, as the genome encodes edited form of the protein.

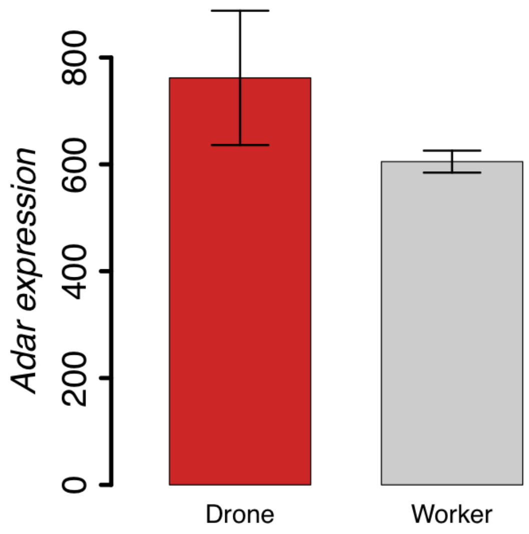


**Figure S11.** ***Adar*** **expression in drones and workers, Related to Figure 4.** Reads count was normalized by DESeq2. Error bars represent standard error of mean. T-test was used to calculate the statistical significance. No significant difference was obtained ($P$ = 0.23).

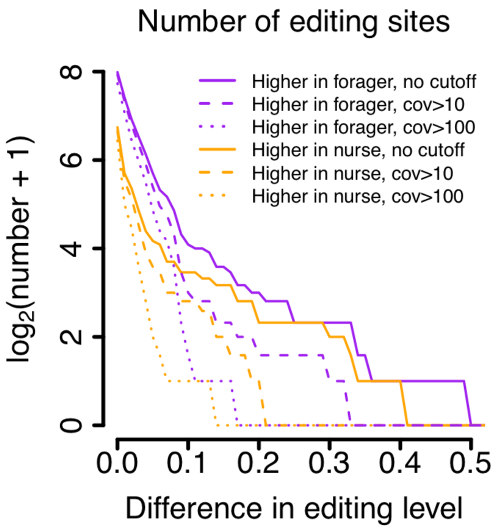


**Figure S12. Comparison of editing levels in two sub-castes of workers, Related to Figure 4.** The numbers of editing sites with levels higher in foragers (purple) and nurses (orange) are shown. Different cutoffs of sequencing coverages were used.

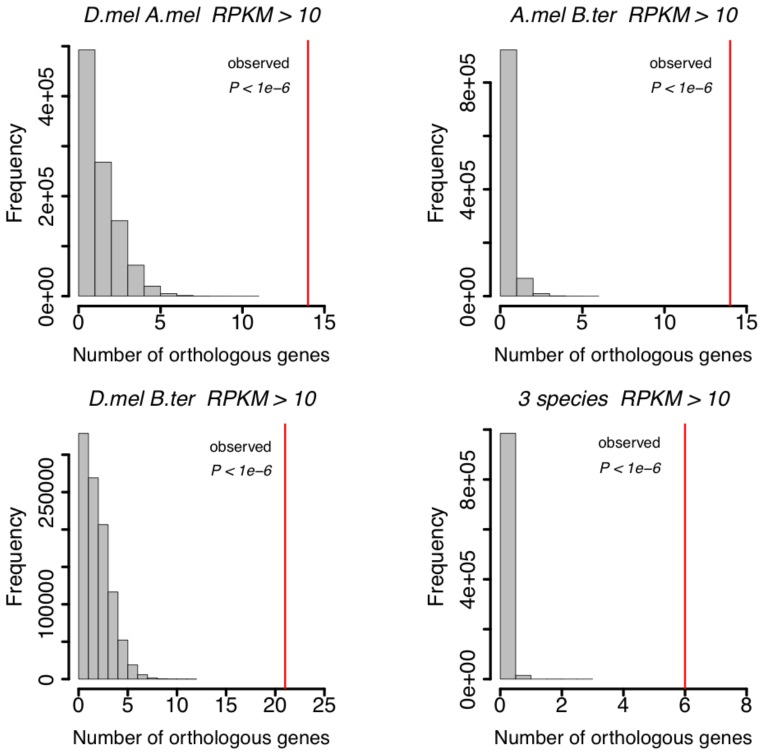


**Figure S13. The observed and expected numbers of orthologous genes with editing in coding regions, Related to Figure 5.** Genes with RPKM > 10 were chosen to calculate the expected numbers.

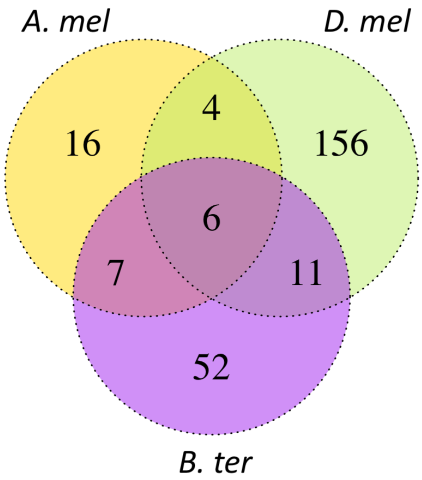


**Figure S14. Venn diagram demonstrating the overlap between orthologous genes with editing sites in coding regions, Related to Figure 5.** Strict criteria for determining sites as edited or not-edited were applied. The details are described in the main text.

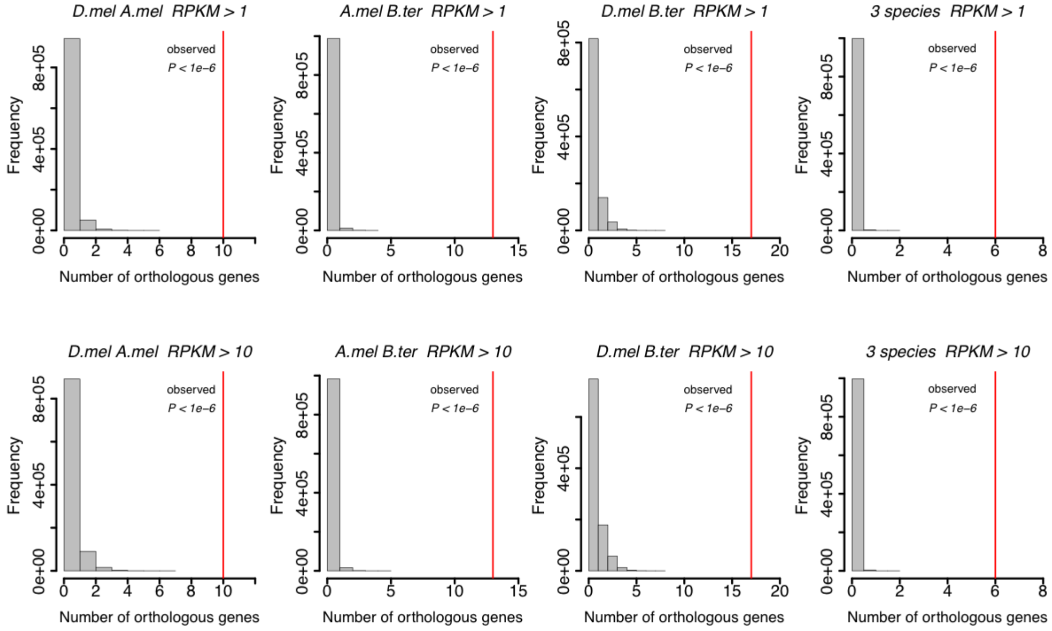


**Figure S15. The observed and expected numbers of orthologous genes with editing in coding regions, Related to Figure 5.** Strict criteria for determining sites as edited or not-edited were applied. The details are described in the main text. Genes with RPKM > 1 or RPKM > 10 were chosen to calculate the expected numbers.

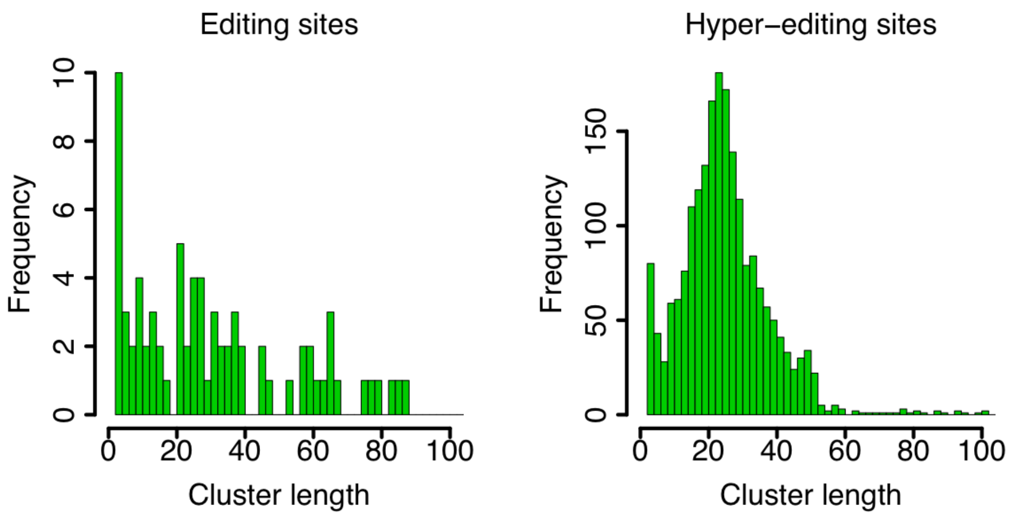


**Figure S16. Histograms showing the cluster length of normal editing sites (left) and hyper-editing sites (right), Related to Figure 1.** Editing sites within 100bps were clustered.

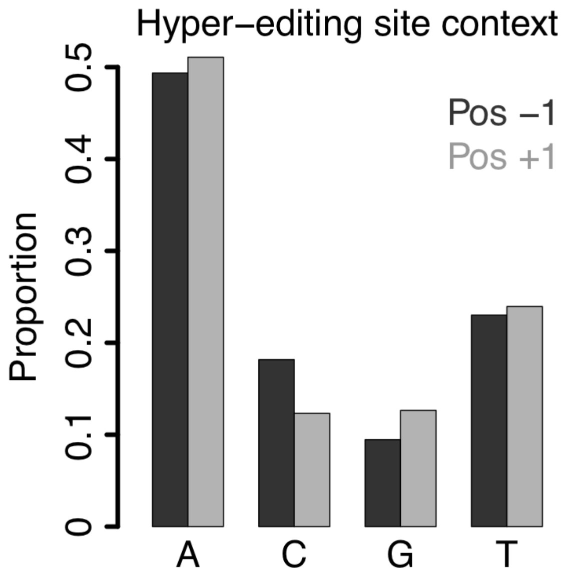


**Figure S17. The nucleotide context of hyper-editing sites, Related to Figure 1.** Pos -1, the upstream nucleotide. Pos +1, the downstream nucleotide.

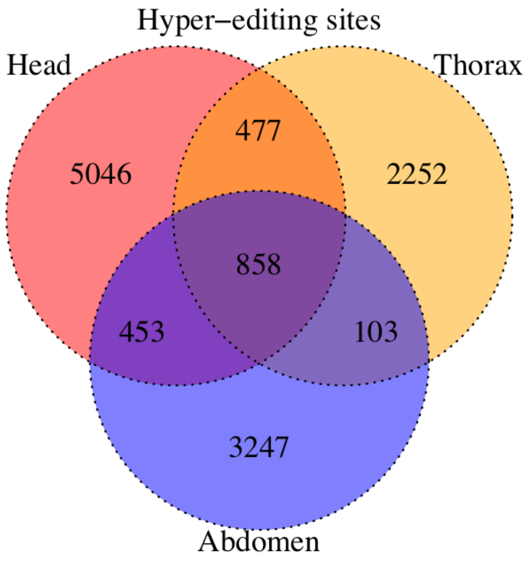


**Figure S18. Venn diagram demonstrating the overlaps of hyper-editing sites in three tissues of honeybee drones, Related to Figure 1.**

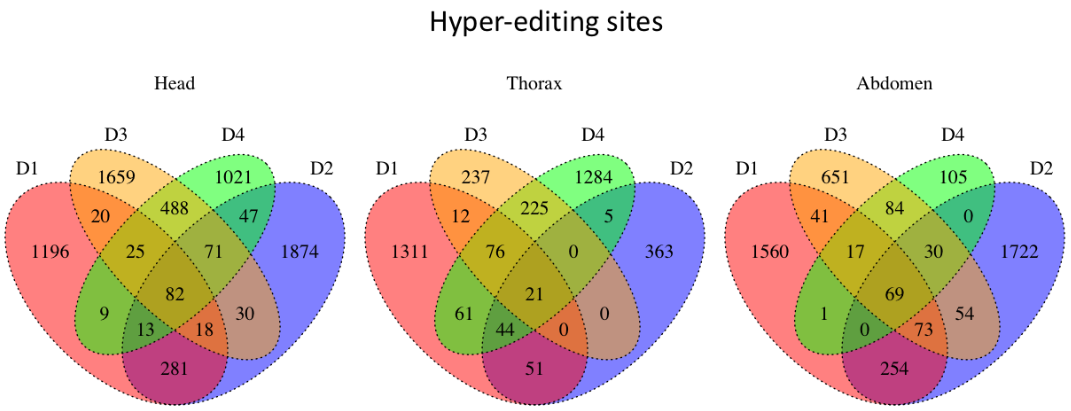


**Figure S19. Venn diagram demonstrating the overlaps of hyper-editing sites in four honeybee drones, Related to Figure 1.**

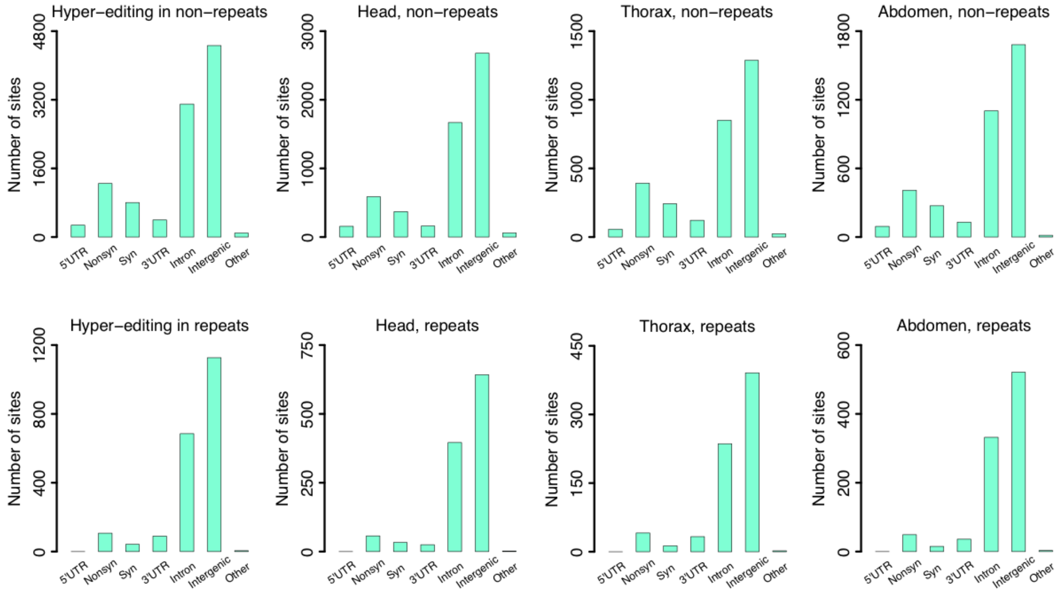


**Figure S20. Annotation of hyper-editing sites for repetitive and non-repetitive regions, separately, Related to Figure 1.** The hyper-editing sites in repeats have lower fractions of nonsynonymous and synonymous sites but have higher fractions of intronic and intergenic sites.

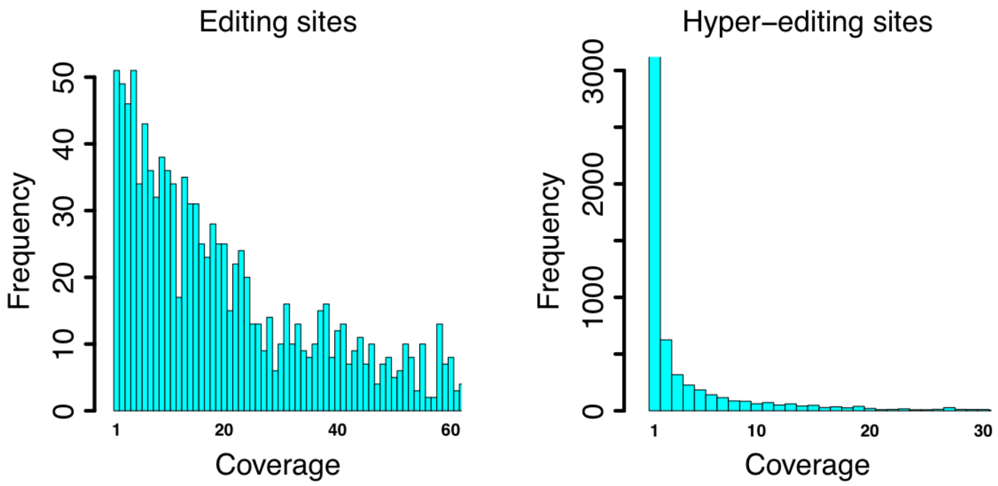


**Figure S21. Histograms of sequencing coverage of normal editing sites (left) and hyper-editing sites (right), Related to Figure 1.** Editing sites in all samples were used.

**Supplemental Tables**

**Table S1. Number of SNPs identified in each individual honeybee, Related to Figure 1.**

| | D1 | D2 | D3 | D4 | All |
|---|---|---|---|---|---|
| 5’UTR | 2,573 | 2,495 | 2,480 | 2,445 | 3,769 |
| Nonsyn | 8,344 | 8,032 | 8,090 | 8,296 | 12,494 |
| Syn | 20,771 | 19,509 | 19,904 | 19,814 | 30,559 |
| 3’UTR | 10,199 | 9,518 | 9,640 | 9,688 | 14,779 |
| Intron | 359,949 | 347,684 | 343,875 | 352,415 | 544,971 |
| Intergenic | 456,956 | 439,499 | 440,563 | 451,767 | 696,412 |
| Other | 157 | 153 | 154 | 159 | 241 |
| Total | 858,949 | 826,890 | 824,706 | 844,584 | 1,303,225 |

**Table S2. Mapping summary of each drone sample, Related to Figure 1.**

| Drones | Tissue | Total reads (M) | Reference genome | | Masked genome | |
|---|---|---|---|---|---|---|
| | | | Uniquely mapped reads (M) | Uniquely mapped reads (%) | Uniquely mapped reads (M) | Uniquely mapped reads (%) |
| D1 | Head | 25.0 | 17.9 | 71.6 | 17.8 | 71.1 |
| | Thorax | 25.8 | 16.8 | 65.1 | 16.1 | 62.4 |
| | Abdomen | 23.5 | 17.2 | 73.2 | 17.2 | 73.1 |
| D2 | Head | 18.8 | 14.2 | 75.3 | 14.1 | 74.8 |
| | Thorax | 30.8 | 18.4 | 59.6 | 17.1 | 55.6 |
| | Abdomen | 28.5 | 21.5 | 75.3 | 21.4 | 75.2 |
| D3 | Head | 37.2 | 12.4 | 33.4 | 13.0 | 34.9 |
| | Thorax | 32.9 | 10.2 | 31.1 | 10.6 | 32.3 |
| | Abdomen | 43.1 | 12.8 | 29.8 | 13.4 | 31.2 |
| D4 | Head | 48.4 | 15.2 | 31.4 | 15.5 | 32.1 |
| | Thorax | 31.9 | 10.4 | 32.6 | 10.7 | 33.4 |
| | Abdomen | 32.9 | 9.2 | 28.1 | 9.5 | 28.8 |

Reference genome: the reads mapped to the reference honeybee genome.

Masked genome: the reads mapped to the genome sequences which have been replaced with the alternative alleles at SNP sites.

**Table S4. Number of editing sites in each sample, Related to Figure 1 and Figure2.**

| Drones | Tissue | Total sites | Nonsyn (*N*) | Syn (*S*) | *N*/*S* | *P* value |
|---|---|---|---|---|---|---|
| D1 | Head | 278 | 72 | 8 | 9.0 | 1.59E-05 |
| | Thorax | 71 | 13 | 4 | 3.3 | 0.610 |
| | Abdomen | 94 | 14 | 1 | 14.0 | 0.0491 |
| D2 | Head | 262 | 73 | 7 | 10.4 | 4.36E-06 |
| | Thorax | 70 | 12 | 5 | 2.4 | 1 |
| | Abdomen | 96 | 14 | 1 | 14.0 | 0.0491 |
| D3 | Head | 177 | 44 | 6 | 7.3 | 0.00318 |
| | Thorax | 40 | 3 | 0 | Inf | 0.558 |
| | Abdomen | 55 | 0 | 0 | 0.0 | 1 |
| D4 | Head | 208 | 59 | 6 | 9.8 | 6.67E-05 |
| | Thorax | 41 | 3 | 1 | 3.0 | 1 |
| | Abdomen | 60 | 5 | 1 | 5.0 | 0.674 |
| All head | | 376 | 108 | 8 | 13.5 | 8.46E-10 |
| All thorax | | 106 | 14 | 5 | 2.8 | 0.807 |
| All abdomen | | 137 | 16 | 1 | 16.0 | 0.0318 |
| All | | 407 | 111 | 9 | 12.3 | 1.02E-09 |

Totally 407 unique editing sites were identified. The numbers of editing sites in each sample are listed respectively. “All head” is the number of sites that appear in at least one head sample among the four individuals. The same goes for “All thorax” and “All abdomen”.

*P* value: The *P* value of the observed *N*/*S* ratios compared to the expected *N*/*S* ratio under neutral evolution calculated from Fisher’s exact tests.

**Table S5. Conserved coding editing sites between honeybee and bumblebee, Related to Figure 3.**

| *Apis mellifera* | | | *Bombus terrestris* | | |
|---|---|---|---|---|---|
| Site | Gene | Editing level | Site | Gene | Editing level |
| Group8.6:728981 | GB40519 | 0.48 | Group16.3:2744556 | XM_003401834 | 0.87 |
| Group8.6:728990 | GB40519 | 0.42 | Group16.3:2744565 | XM_003401834 | 0.41 |
| GroupUn131:18550 | GB46982 | 0.31 | Group10.1:6435961 | XM_003398309 | 0.93 |
| GroupUn131:18489 | GB46982 | 0.79 | Group10.1:6436022 | XM_003398309 | 0.97 |
| GroupUn131:18488 | GB46982 | 0.81 | Group10.1:6436023 | XM_003398309 | 0.98 |
| Group1.29:1345934 | GB47508 | 0.50 | Group1.8:1831958 | XM_003393355 | 0.88 |
| Group1.29:1345950 | GB47508 | 0.34 | Group1.8:1831942 | XM_003393355 | 0.79 |
| Group15.19:753243 | GB50090 | 0.42 | Group15.5:2877069 | XM_003401089 | 0.45 |
| Group1.29:1555391 | GB47375 | 0.70 | Group1.8:1578895 | XM_003393369 | 0.63 |

**Table S6. Mapping summary of each worker sample, Related to Figure 4.**

| Sample | Total reads (M) | Uniquely mapped reads (M) | Uniquely mapped reads (%) |
|---|---|---|---|
| SRR445999 | 213.8 | 118.2 | 55.3 |
| SRR446000 | 295.4 | 203.8 | 69.0 |
| SRR446001 | 152.0 | 95.2 | 62.6 |
| SRR446002 | 154.5 | 87.9 | 56.9 |
| SRR446003 | 138.6 | 89.2 | 64.4 |
| SRR446004 | 226.0 | 149.9 | 66.3 |
| Nurse pool | 1180.3 | 744.2 | 63.0 |
| SRR446005 | 178.9 | 97.5 | 54.5 |
| SRR446006 | 193.1 | 130.3 | 67.5 |
| SRR446007 | 121.2 | 72.8 | 60.1 |
| SRR446008 | 164.6 | 100.2 | 60.9 |
| SRR446009 | 179.7 | 119.4 | 66.5 |
| SRR446010 | 139.4 | 84.5 | 60.6 |
| Forager pool | 976.8 | 604.6 | 61.9 |

**Table S9. Shared genes with CDS editing across three species, Related to Figure 5.**

| *D. melanogaster* | *A. mellifera* | *B. terrestris* |
|---|---|---|
| FBgn0086372 | GB43906 | XM_003403010 |
| FBgn0263354 | GB48155 | XM_003396523 |
| FBgn0263111 | GB51897 | XM_003394356 |
| FBgn0035538 | GB54467 | XM_003395885 |
| FBgn0004242 | GB54827 | XM_003402654 |
| FBgn0262483 | GB55567 | XM_003402632 |

The gene IDs of *B. terrestris* are retrieved from the bumblebee study (Porath et al. 2019) according to the editing site.

**Table S10. Number of hyper-editing sites in each sample, Related to Figure 1.**

| Drones | Tissue | Total sites | 5'UTR | Nonsyn | Syn | 3'UTR | Intron | Intergenic | other |
|---|---|---|---|---|---|---|---|---|---|
| D1 | Head | 1,644 | 7 | 86 | 50 | 72 | 509 | 915 | 5 |
| | Thorax | 1,576 | 32 | 182 | 100 | 58 | 387 | 801 | 16 |
| | Abdomen | 2,015 | 29 | 215 | 114 | 65 | 607 | 977 | 8 |
| D2 | Head | 2,416 | 110 | 290 | 170 | 37 | 660 | 1,110 | 39 |
| | Thorax | 484 | 19 | 54 | 44 | 46 | 143 | 172 | 6 |
| | Abdomen | 2,202 | 64 | 221 | 153 | 68 | 730 | 954 | 12 |
| D3 | Head | 2,393 | 37 | 196 | 135 | 52 | 817 | 1,143 | 13 |
| | Thorax | 571 | 0 | 44 | 32 | 9 | 168 | 318 | 0 |
| | Abdomen | 1,019 | 6 | 57 | 49 | 17 | 291 | 598 | 1 |
| D4 | Head | 1,756 | 11 | 145 | 92 | 29 | 495 | 977 | 7 |
| | Thorax | 1,716 | 18 | 189 | 102 | 42 | 602 | 755 | 8 |
| | Abdomen | 306 | 0 | 18 | 8 | 17 | 123 | 138 | 2 |
| Pool | | 12,436 | 276 | 1,355 | 845 | 488 | 3,784 | 5,591 | 97 |

### Transparent Methods

#### Honeybee collection

Drones were raised in a colony of *Apis mellifera* by professional beekeepers in Jie Wu's Lab, Chinese Academy of Agricultural Sciences. For each male drone adult (haploid), the head and thorax were separated with surgical scissors and the testis was dissected. Residual abdomen tissues were preserved for DNA extraction. All samples were flash-frozen in liquid nitrogen and stored at -80°C for further procedures.

#### RNA extraction and mRNA-Seq of four drone individuals

Four drone individuals were selected for mRNA-Seq. Total RNA was extracted from head, thorax, and testis from each individual, separately, using TRIzol reagent (Thermo Fisher). For two individuals (1 and 2), Poly(A)+ mRNAs were selected on oligo-dT25 DynaBeads (Thermo Fisher), while for the other two (3 and 4) RNAs were treated with Ribo-Zero Gold rRNA Removal Kit (Illumina) to remove the rRNA-derived fragments. All these purified RNAs were fragmented. The 40-80 nt fragments were purified from 15% TBE-Urea gels for deep sequencing, and subject to 3'-dephosphorylation with T4 Polynucleotide Kinase (NEB), 3'-ligation, 5'-phosphorylation with T4 Polynucleotide Kinase (NEB) and ATP, 5'-ligation, and reverse-transcription into cDNA with SuperScript™ III Reverse Transcriptase (Thermo Fisher). The cDNAs were PCR-amplified and size-selected in 20% TBE gels for fragments in correct ranges. Purified products were prepared for quality tests (Fragment Analyzer, Agilent Technologies) and sequencing (Illumina HiSeq-2500 sequencer; run type: single-end; read length: 50 nt).

#### Genomic sequencing of four drone individuals

To efficiently exclude SNPs from RNA editing sites, genomic DNA from abdomen tissue of each individual drone separately was extracted using the Genomic DNA Extraction Kit (TIANGEN) following manufacturer's instructions. The library preparation and sequencing were performed in Biomedical Pioneering Innovation Center, Peking University (Illumina HiSeq-2500 sequencer; 100bp paired-end reads).

#### Identification of SNPs in four drones

Honeybee reference genome sequence (*A. mel* 4.5) was downloaded from BeeBase (http://hymenopteragenome.org/beebase/). For each of the four individual drones, we mapped the DNA-Seq reads to the reference genome with BWA v0.7.4 (Li and Durbin 2009). PCR duplicates were removed using Picard v1.119, and SNPs were called using SAMtools mpileup (Li 2011) with default parameters.

**Identification of variation sites in four drones**

To identify reliable A-to-I RNA editing events, we employed two different aligners STAR (2.4.2a) (Dobin et al. 2013) and BWA (Li and Durbin 2009) to map the RNA-Seq reads to the reference genome (*A. mel* 4.5). For each BAM sequence alignment file, we extracted all the alignments with mapping quality ≥ 10 using SAMtools 1.3.1 (Li 2011). The mismatches between RNA-Seq reads and reference genome were extracted using Sam2Tsv (Pierre 2015). Bases with mapping quality lower than 30, soft clipping bases, bases at the 10bp of reads' ends and mismatches in repeat regions were discarded. Only mismatch sites supported by both STAR and BWA were retained.

Next, we produced for each individual drone its own version of the genome, by substituting the SNP sites found in this individual to the reference genome. For example, in individual drone1, 858,949 SNPs were identified, and the reference genome (*A. mel* 4.5) was modified by replacing the reference alleles with the alternative alleles at all these SNP sites of drone1. The modified genome was regarded as the masked genome for drone1. Note that drone individuals are haploids and there are no allele-specific SNPs. Thus, the masked genome is the entire haploid genome sequence of each individual. We then mapped again the RNA-Seq reads of drone 1 to its masked genome, using both STAR and BWA as before. The same pipeline was applied to drones 2, 3, and 4.

Ideally, one would need to map the reads only to the masked genome (presumably, the true genome of each individual drone). However, our SNP detection is imperfect, and false positive SNPs identification could lead to RNA-DNA variations against the masked genome. Thus, to be conservative, we retained only variation sites supported by mapping to both the reference genome and the masked genomes. These RNA-DNA variations are unlikely to include many variations due to genomic polymorphisms, and should be enriched in RNA level alterations. We further discarded variations occurring in less than three out of twelve samples (head, thorax, and abdomen of four honeybee individuals), resulting in 1,742 candidate variation sites. Of these, 1417 (81.3%) were A-to-G.

**Defining A-to-I RNA editing sites**

Finally, we pooled the reads of the four drones for each of the tissues, and calculated the probability $P_k(E_0)$ that the mismatches at position $E_0$ (each of the 1,742 sites found above) observed in tissue $k$ ($k$ = head, thorax, or abdomen) can be explained by sequencing error (binomial test, with A-to-G error rate $\varepsilon = 0.00167$ (Duan et al. 2017), followed by Benjamini-Hochberg multiple testing correction with FDR = 0.05 (Benjamini and Hochberg 1995)). The multiple testing correction took into account all ~230 Mbp of the honeybee genome (excluding "N"s), at which the above variations may have been detected. Pooled data was analyzed separately for each tissue.

Editing level was estimated as G/(G+A), in which G is the alternative allele count and A is the reference allele count.

For the downloaded deep-sequencing data of brains of honeybee workers (Herb et al. 2012), we mapped the reads with STAR and called variants with SAMtools mpileup (Li 2011) with default parameters as we mentioned above. We directly retrieved the reads counts on the candidate editing sites and looked at the editing levels. Alignments with mapping quality ≥ 10 were maintained and no other filters were applied. Editing levels was compared between different castes or sub-castes using Fisher's exact test. Editing levels in other species (e.g. sites in genes *Shab* and *qvr*), were evaluated based on the datasets described below. We mapped the sequencing reads of each species to the reference genome using STAR (Dobin et al. 2013). The variants were called with SAMtools mpileup (Li 2011). The genomic coordinates were transferred from *D. melanogaster* to the other fly species by using liftOver chain downloaded from UCSC Genome Browser (http://genome.ucsc.edu/).

**Annotation of A-to-I RNA editing sites**

We used the software SnpEff version 4.3 (Cingolani et al. 2012b) for functional annotation of the editing sites. The canonical transcript of each gene, as defined by the software, was chosen for annotation.

**Expected *N*/*S* ratio**

To find the expected *N*/*S* ratio under neutrality, we replaced all adenosines in all coding regions with guanosines, one at a time, and used the software SnpEff (Cingolani et al. 2012a), (version 4.3, parameters "-ud -canon -v amel_OGSv3.2) to determine how many of these substitutions would cause a nonsynonymous or synonymous change. In total, 4,492,737 nonsynonymous and 1,986,130 synonymous substitutions were obtained, and therefore the expected *N*/*S* ratio was 2.26. To verify that this ratio is not biased by editing being limited to a small set of genes, we have repeated the calculation focusing only on adenosines in genes harboring editing sites. Here we have found 68,220 nonsynonymous and 30,150 synonymous substitutions were obtained, and therefore the expected *N*/*S* ratio was very similar, 2.26. We therefore used the ratio 2.26 is all further analyses.

**Conservation analysis**

Orthologous genes (between *D. melanogaster* and honeybee, or between honeybee and bumblebee) were defined as reciprocal best hits of pairwise BLASTP (Camacho et al. 2009). The protein sequences of the orthologous genes were then aligned with the clustalw program (Thompson et al. 1994), and the alignments of RNA CDS were achieved by the tranalign program (Rice et al. 2000) based on the corresponding protein alignments. We used *D. melanogaster* reference genome version FlyBase_r6.04. For the bumblebee genome we used NCBI reference sequence version Bter_1.0. NCBI RefSeq annotations for genes and coding regions were downloaded from the BeeBase site (http://hymenopteragenome.org/beebase) on 7 May, 2014.

To calculate the expected numbers of genes sharing CDS editing at different positions within the same gene we first determined the set of expressed genes in in each of the three species as those genes with RPKM (reads per kilobase per million mapped reads) values exceeding a given cutoff. Then, we have randomly chosen 53 honeybee genes, 101 bumblebee genes and 312 fly genes (the numbers of genes harboring editing sites in CDS per species, excluding sites conserved in any two or three species), and looked for the number of orthologous gene groups reoccurring in any two of these sets or in all three of them. For each expression cutoff, the distribution of the expected overlap was calculated using 1,000,000 such random choices, and the p-value was calculated comparing the actual overlap to this distribution.

**mRNA secondary structure prediction**

We folded the mRNAs of honeybee with RNALfold (Lorenz et al. 2011). Local structures with Z score < -3 were defined as stable hairpins.

**PCA Analysis**

The PCA analysis of editing levels and gene expression levels was performed by function "princomp()" in R. The first two principal components (PC1 and PC2) were used to plot the graph.

**Supplemental Note**

**The hyper-editing pipeline detects many sites in lowly expressed regions**

We used the hyper editing script (Porath et al. 2014) with parameters suitable for 50bp reads "0.05 0.8 30 0.6 0.1 0.8 0.2" (Porath et al. 2014) to identify identified densely edited clusters, with a total of 12,436 unique sites. Of these, only 2,056 (17%) are located in repetitive regions. This fraction is higher than the one found by our main procedure, but still much lower than the one observed in most other species analyzed. A partial explanation is the rather short reads, preventing reliable alignment of heavily edited reads.

Hyper-editing sites are clustered. Clustering together editing sites (distance < 100 bps) we find that hyper-editing sites tend to cluster in big clusters (larger than the ones required for their detection) (Figure S16). Notably, half of regular editing sites are also clustered.

The sequence context of these hyper-editing sites is similar to the pattern observed in bumblebees (Porath et al. 2019), where the upstream nucleotide shows a preference of A > T > C > G and the downstream nucleotide exhibits A > T > G > C (Figure S17). In each sample, 306~2,416 hyper editing sites were detected in different tissues of four honeybee individuals (Table S10). Heads bear the greatest numbers of hyper editing sites than other tissues (Figure S18), although the sites from heads of different individuals do not show much overlap (Figure S19).

Compared to the editing sites identified by our regular pipeline, those hyper-editing sites have a remarkably higher fraction in intergenic and intron regions (Figure S20). In fact, only 52 of these 12,436 sites overlap with the 407 editing sites identified by traditional pipeline, and 45 of the overlapped sites are located in intergenic or intronic region. This result agrees with the finding that hyper-edited regions are often lowly expressed (Porath et al. 2017), and corroborated by the sequencing coverage spectrum which for hyper-editing sites is skewed to lower depths compared to normal editing sites (Figure S21). None of the hyper-editing sites overlap with conserved or task-related editing sites in the bumblebee, which further suggests that hyper-edited regions are weakly expressed and likely to be non-conserved.

### Supplemental References (Related to Supplemental Figures, Supplemental Tables, and Transparent Methods)


Benjamini Y, Hochberg Y. 1995. Controlling the False Discovery Rate - a Practical and Powerful Approach to Multiple Testing. *Journal of the Royal Statistical Society Series B-Methodological* **57**: 289-300.

Camacho C, Coulouris G, Avagyan V, Ma N, Papadopoulos J, Bealer K, Madden TL. 2009. BLAST+: architecture and applications. *BMC Bioinformatics* **10**: 421.

Cingolani P, Platts A, Wang le L, Coon M, Nguyen T, Wang L, Land SJ, Lu X, Ruden DM. 2012a. A program for annotating and predicting the effects of single nucleotide polymorphisms, SnpEff: SNPs in the genome of Drosophila melanogaster strain w1118; iso-2; iso-3. *Fly (Austin)* **6**: 80-92.

Cingolani P, Platts A, Wang LL, Coon M, Nguyen T, Wang L, Land SJ, Lu X, Ruden DM. 2012b. A program for annotating and predicting the effects of single nucleotide polymorphisms, SnpEff. *Fly* **6**: 80-92.

Dobin A, Davis CA, Schlesinger F, Drenkow J, Zaleski C, Jha S, Batut P, Chaisson M, Gingeras TR. 2013. STAR: ultrafast universal RNA-seq aligner. *Bioinformatics* **29**: 15-21.

Duan Y, Dou S, Luo S, Zhang H, Lu J. 2017. Adaptation of A-to-I RNA editing in Drosophila. *PLoS Genet* **13**: e1006648.

Herb BR, Wolschin F, Hansen KD, Aryee MJ, Langmead B, Irizarry R, Amdam GV, Feinberg AP. 2012. Reversible switching between epigenetic states in honeybee behavioral subcastes. *Nat Neurosci* **15**: 1371-1373.

Li H. 2011. A statistical framework for SNP calling, mutation discovery, association mapping and population genetical parameter estimation from sequencing data. *Bioinformatics* **27**: 2987-2993.

Li H, Durbin R. 2009. Fast and accurate short read alignment with Burrows-Wheeler transform. *Bioinformatics* **25**: 1754-1760.

Lorenz R, Bernhart SH, Honer Zu Siederdissen C, Tafer H, Flamm C, Stadler PF, Hofacker IL. 2011. ViennaRNA Package 2.0. *Algorithms for molecular biology : AMB* **6**: 26.

Pierre L. 2015. *JVarkit: java-based utilities for Bioinformatics*.

Porath HT, Carmi S, Levanon EY. 2014. A genome-wide map of hyper-edited RNA reveals numerous new sites. *Nature Communications* **5**.

Porath HT, Hazan E, Shpigler H, Cohen M, Band M, Ben-Shahar Y, Levanon EY, Eisenberg E, Bloch G. 2019. RNA editing is abundant and correlates with task performance in a social bumblebee. *Nature Communications* **10**.

Porath HT, Knisbacher BA, Eisenberg E, Levanon EY. 2017. Massive A-to-I RNA editing is common across the Metazoa and correlates with dsRNA abundance. *Genome Biol* **18**: 185.

Rice P, Longden I, Bleasby A. 2000. EMBOSS: the European Molecular Biology Open Software Suite. *Trends Genet* **16**: 276-277.

Thompson JD, Higgins DG, Gibson TJ. 1994. CLUSTAL W: improving the sensitivity of progressive multiple sequence alignment through sequence weighting, position-specific gap penalties and weight matrix choice. *Nucleic Acids Res* **22**: 4673-4680.
